# A TAK1-Driven NLRP1 Inflammasome Pathway Revealed by Phosphatase-Targeting Environmental Toxins

**DOI:** 10.64898/2026.01.23.701233

**Authors:** Margaux Paradis, Léana Gorse, Leandro Silva Da Costa, Rae Chua, Geraldine Aberola, Arnaud Metais, Ludovic Batistic, Thomas Benoist, Anthony David, Emeline Pagès, Andrea Gomes, Caïo Bomfim, Léa Ravon-Katossky, Bastien Suire, Léa Fromont, Ron Siaden-Ortega, David Péricat, Céline Cougoule, Laurent Boyer, Nicolas Gaudenzio, Raoul Mazars, Virginie Stevenin, Franklin Lei Zhong, David Olagnier, Etienne Meunier

## Abstract

Human NLRP1 is highly sensitive to phosphorylation-dependent activation, indicating a need for tight phosphatase control to prevent excessive inflammasome activity. In this study, we identified the phosphatases PP1 and PP2A as negative regulators of the NLRP1 inflammasome in human keratinocytes. Accordingly, exposure to the environmental toxins Dinophysistoxin, Okadaic acid, and Cantharidin, which inhibit PP1 and PP2A, triggered NLRP1 inflammasome activation. Notably, this toxin-induced activation process relied on hyperactivation of the MAP3 kinase TAK1. Mechanistically, both TAK1 and its downstream effectors p38 kinases phosphorylated and activated the NLRP1 inflammasome. Further studies underscored that TAK1 also contributed to the NLRP1 inflammasome activation during double-stranded RNA stimulation and viral infection. Finally, human native skins exposed to environmental toxins underlined the role of the PP1/PP2A-regulated TAK1/p38 axis in the development of skin dermatitis. Thus, these findings reveal a novel pathway of phosphorylation-driven NLRP1 activation and expand our understanding of its regulation in epithelial immunity.

**One Sentence Summary:** PP1/PP2A phosphatases restrict TAK1-driven NLRP1 inflammasome response.

Graphical abstract
Schematic representation of the phosphorylation-driven hNLRP1 inflammasome response upon exposure to environmental toxins and to dsRNA/viral infection.

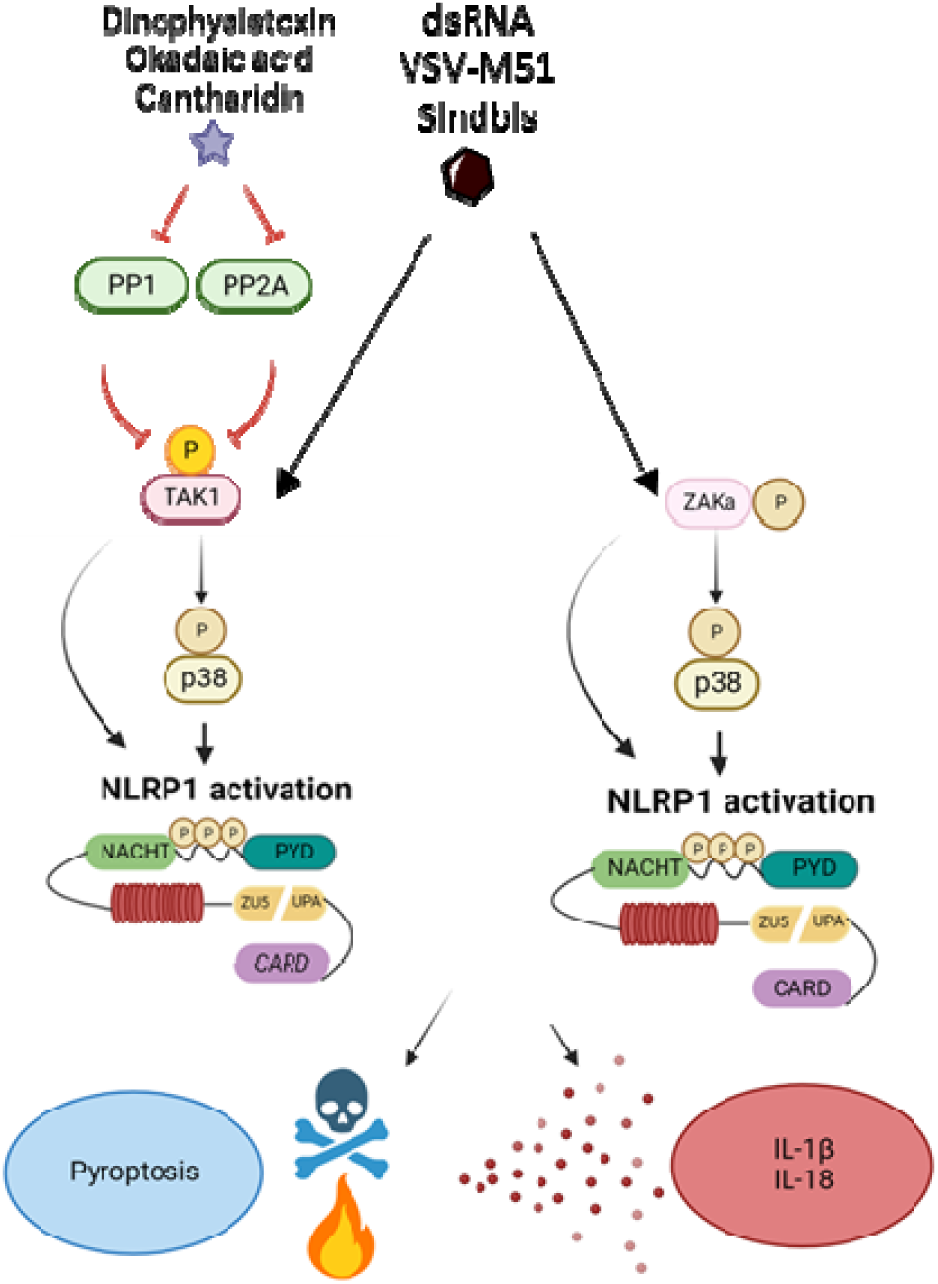

## INTRODUCTION

To cope with an environment filled with living microorganisms and (non)-biological threats such as tissue injury, UV radiation, or chemical exposure, the innate immune system has evolved a diverse repertoire of pattern-recognition receptors (PRRs) (*1*, *2*). These receptors, expressed across numerous cell types, play a central role in coordinating both inflammatory and non-inflammatory defenses (*2*, *3*). Among PRRs, the inflammasome-forming sensors constitute a specialized family capable of coupling the detection of danger signals to both cytokine release and programmed inflammatory cell death (*1*, *2*).

Inflammasomes are cytosolic signaling complexes that, upon activation by pathogens or cellular stress, induce pyroptotic cell death and the secretion of the pro-inflammatory cytokines IL-1β and IL-18 (*1*). This process requires the protease caspase-1, which cleaves and activates IL-1β/IL-18 and processes gasdermin D to form membrane pores, ultimately culminating in pyroptosis and Ninjurin-1–mediated cell lysis (*1*, *4–8*). Although inflammasome activation is essential for antimicrobial defense, dysregulation can drive chronic autoinflammation and severe tissue injury (*2*).

Human NLRP1 is distinctive among inflammasome sensors because of its robust expression in epithelial tissues, including keratinocytes, the airway epithelium, and the cornea (*9*). A hallmark of NLRP1 activation is the release of its C-terminal CARD-containing fragment following proteasome-dependent degradation of its N-terminal region, a mechanism termed “functional degradation.” The liberated CARD fragment then oligomerizes to recruit and activate caspase-1 (*10*).

Recent studies have established NLRP1 as a versatile detector of redox and proteotoxic stress (*11–13*), as well as viral proteases from picornaviruses and coronaviruses (*14–16*). Moreover, NLRP1 is activated by ZAKα- and p38-driven phosphorylation in response to ribotoxic stress response triggered by viral double stranded RNA, bacterial toxins, and environmental insults (*9*, *17–22*).

Given the high sensitivity of hNLRP1 to phosphorylation-induced activation, we hypothesized that cellular phosphatases might act as critical brakes to prevent uncontrolled NLRP1 signaling (*23–25*). In this study, we identified the serine/threonine phosphatases PP1 and PP2A (*23–25*) as key negative regulators of the NLRP1 inflammasome in human keratinocytes. Consistent with this, exposure to environmental toxins such as dinophysistoxin, okadaic acid, and cantharidin, all potent inhibitors of PP1 and PP2A, triggered robust NLRP1 activation. We found that this response was driven by hyperactivation of the MAP3K TAK1. Mechanistically, both TAK1 and its downstream effectors, the p38 MAP kinases, phosphorylated and activated hNLRP1. Furthermore, we also unveiled that TAK1 contributed to NLRP1 activation in response to double stranded RNA and viral infection, suggesting that this pathway extends beyond phosphatase inhibition. Finally, experiments using human skin explant models exposed to environmental toxins demonstrated that the PP1/PP2A-TAK1-p38 axis plays a central role in promoting toxin-induced dermatitis. Together, our findings uncover a previously unrecognized mechanism in which inhibition of PP1/PP2A phosphatases unleashes a TAK1/p38-driven phosphorylation cascade that activates the NLRP1 inflammasome.

## RESULTS

### A pharmacological screen identifies PP1/PP2A phosphatase inhibitors and toxins as inducers of the human NLRP1 inflammasome

Given the high sensitivity of NLRP1 to phosphorylation-dependent activation, we hypothesized that multiple phosphatases might tightly regulate the phosphorylation/dephosphorylation dynamics governing hNLRP1 inflammasome activity. To explore this, we performed a pharmacological screen coupled with fluorescence microscopy in HEK293 reporter cells expressing hNLRP1 and ASC-GFP (HEK293 ASC-GFP/NLRP1). Cells were exposed to a small library of phosphatase inhibitors, and inflammasome activation was quantified 8 hours later by assessing ASC-GFP speck formation (Fig. 1A, B). Among all tested inhibitors, ten compounds induced detectable hNLRP1 inflammasome assembly (Fig. 1A, B and Fig. S1A).

**Figure 1.**
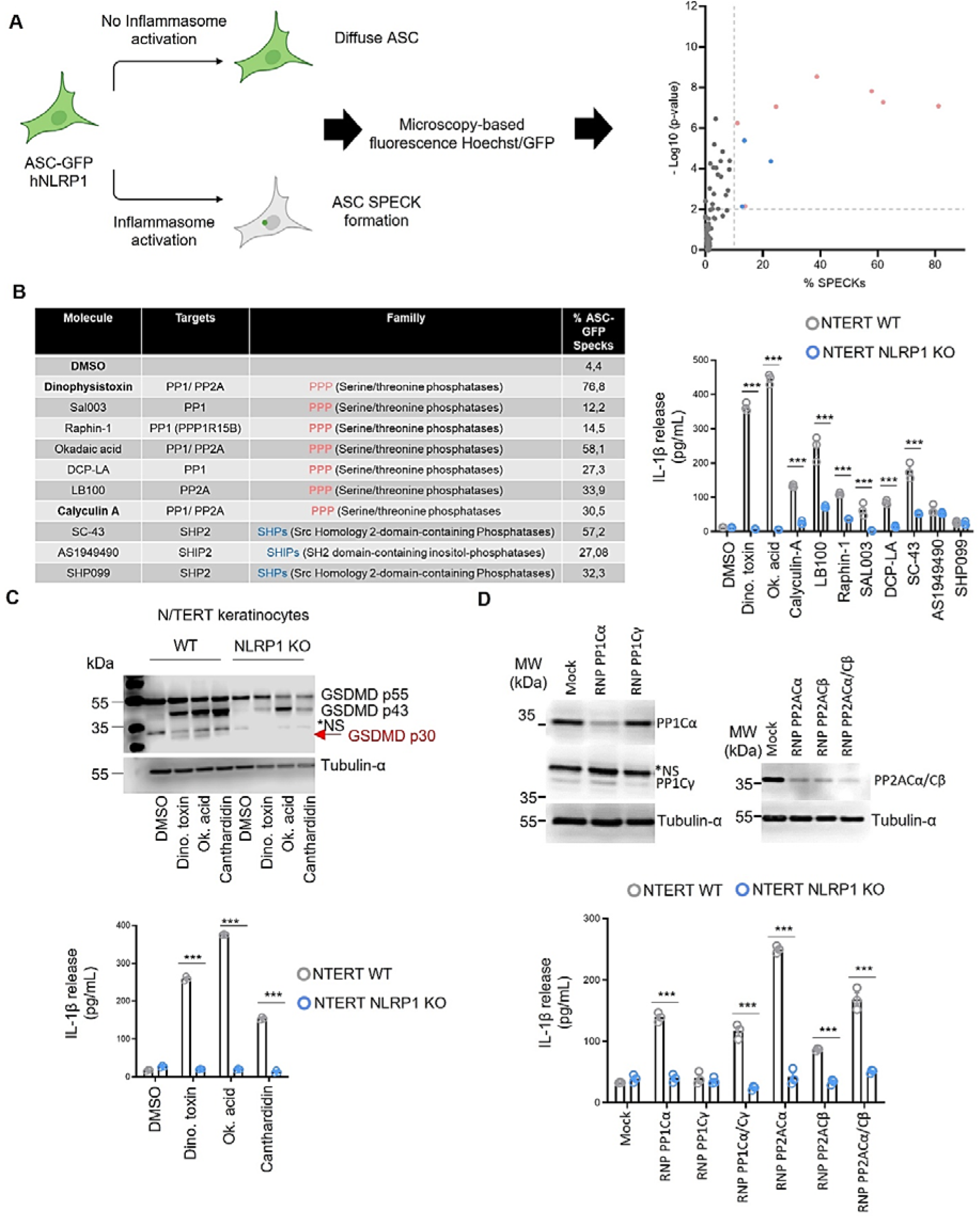
A pharmacological screen identifies PP1/PP2A phosphatase inhibitors and toxins as inducers of the human NLRP1 inflammasome. **A, B.** Screening methodology, associated quantifications of ASC-GFP specks in HEK293T cells individually expressing or not NLRP1 and IL-1β release in NTERT-keratinocytes (WT and NLRP1KO) exposed to 1µM of compounds from the phosphatase screening library for 8 h. ASC-GFP (green) pictures were taken with an EVOS7000 after adding Hoechst (nuclei staining) directly in medium. The percentage of ASC complex was performed by determining the ratios between cells positive for ASC speckles and the total of cell nuclei (Hoechst) by automatic fluorescence microscopy. At least ten fields from each experiment were analyzed. Values are expressed as mean ± SEM. ***P ≤ 0.0001, one-way ANOVA. Graphs show one experiment performed in triplicates at least three times. **C.** Immunoblotting of GSDMD and analysis of the subsequent IL-1β release in WT and NLRP1-deficient NTERT-keratinocytes after 8 h exposure to Dinophysis toxin (Dino. toxin, 100nM), Okadaic acid (Ok. Acid, 250nM) and Cantharidin (Canth. 5µM). Immunoblots show combined lysates+supernatants from one experiment performed at least three times. For cytokine release, ***P ≤ 0.0001, two-way ANOVA with multiple comparisons. Values are expressed as mean ± SEM. Graphs show one experiment performed in triplicates at least three times. **D.** Immunoblotting of PP1 and PP2A catalytic subunits (PP1Cα or PP1Cγ and PP2Acα or PP2Acβ) and analysis of the subsequent IL-1β release in WT and NLRP1-deficient NTERT-keratinocytes 24 hours after CRISPR-Cas9 (RNP)-mediated PP1/PP2A catalytic subunit invalidation. For cytokine release, ***P ≤ 0.0001, two-way ANOVA with multiple comparisons. Values are expressed as mean ± SEM. Graphs show one experiment performed in triplicates at least three times.

To validate these findings in a more physiological model, we treated WT and NLRP1-deficient human TERT keratinocytes with the ten candidate compounds and measured IL-1β release as a readout of inflammasome activation (Fig. 1B). Eight of the ten inhibitors reproducibly induced NLRP1-dependent IL-1β secretion. Analysis of their described molecular targets revealed that seven compounds inhibit the PP1/PP2A family of serine/threonine phosphatases, whereas the remaining three target the tyrosine phosphatase SHP2 (Fig. 1A, B). Although the potential involvement of tyrosine phosphatases in NLRP1 regulation is intriguing, the strong enrichment of PP1/PP2A-targeting compounds prompted us to focus on these phosphatases.

Among the identified PP1/PP2A-targeting compounds, were the environmental toxins dinophysistoxin-1, okadaic acid, and calyculin A (*26*), and our bibliographic study added the blister beetle toxin cantharidin (*27*), all able to induce skin dermatitis (*28*, *29*). In this context, using a panel of PP1/PP2A-targeting toxins, including the algal toxins dinophysistoxin-1, okadaic acid, and calyculin A, and the blister beetle toxin cantharidin, we confirmed that all these molecules robustly activated the NLRP1 inflammasome in human keratinocytes and in reporter cells, as evidenced by ASC speck assembly, IL-1β release, and gasdermin-D processing (Fig. 1A–C, Fig. S1B-D).

Finally, in order to strengthen these observations, we genetically disrupted several catalytic subunits of PP1 and PP2A (PP1 Cα/Cγ and PP2A Cα/Cβ) in WT and NLRP1-KO TERT keratinocytes using CRISPR–Cas9 (Fig. 1D) (*23–25*). Loss of PP1 Cα or PP2A Cα/Cβ were sufficient to induce IL-1β release in an NLRP1-dependent manner, indicating that multiple PP1/PP2A phosphatase subunits could contribute to restraining NLRP1 inflammasome activation.

Altogether, our results identify PP1 and PP2A phosphatases as key toxin-sensitive negative regulators of the NLRP1 inflammasome in human keratinocytes.

### PP1/PP2A-targeting toxins activate the NLRP1 inflammasome in a ZAK**α**-independent manner

The robust NLRP1-dependent response induced by PP1/PP2A inhibition suggested that PP1/PP2A phosphatases may normally restrain one or more kinases responsible for hNLRP1 phosphorylation and activation. hNLRP1 phosphorylation occurs within its disordered region (DR, aa 86–275), which contains multiple serine and threonine residues highly sensitive to phosphorylation, an essential step for phosphorylation-driven NLRP1 inflammasome activation (Fig. 2A) (*17*, *18*, *20–22*). To investigate whether PP1/PP2A-targeting toxins promote phosphorylation of this region, we used HEK293 cells expressing the hNLRP1 DR fused to GFP (hNLRP1-DR–GFP) and assessed its phosphorylation status following toxin or compound treatment (Fig. 2B). Anisomycin, a known inducer of hNLRP1 phosphorylation via the ribotoxic stress response, served as a positive control.

**Figure 2.**
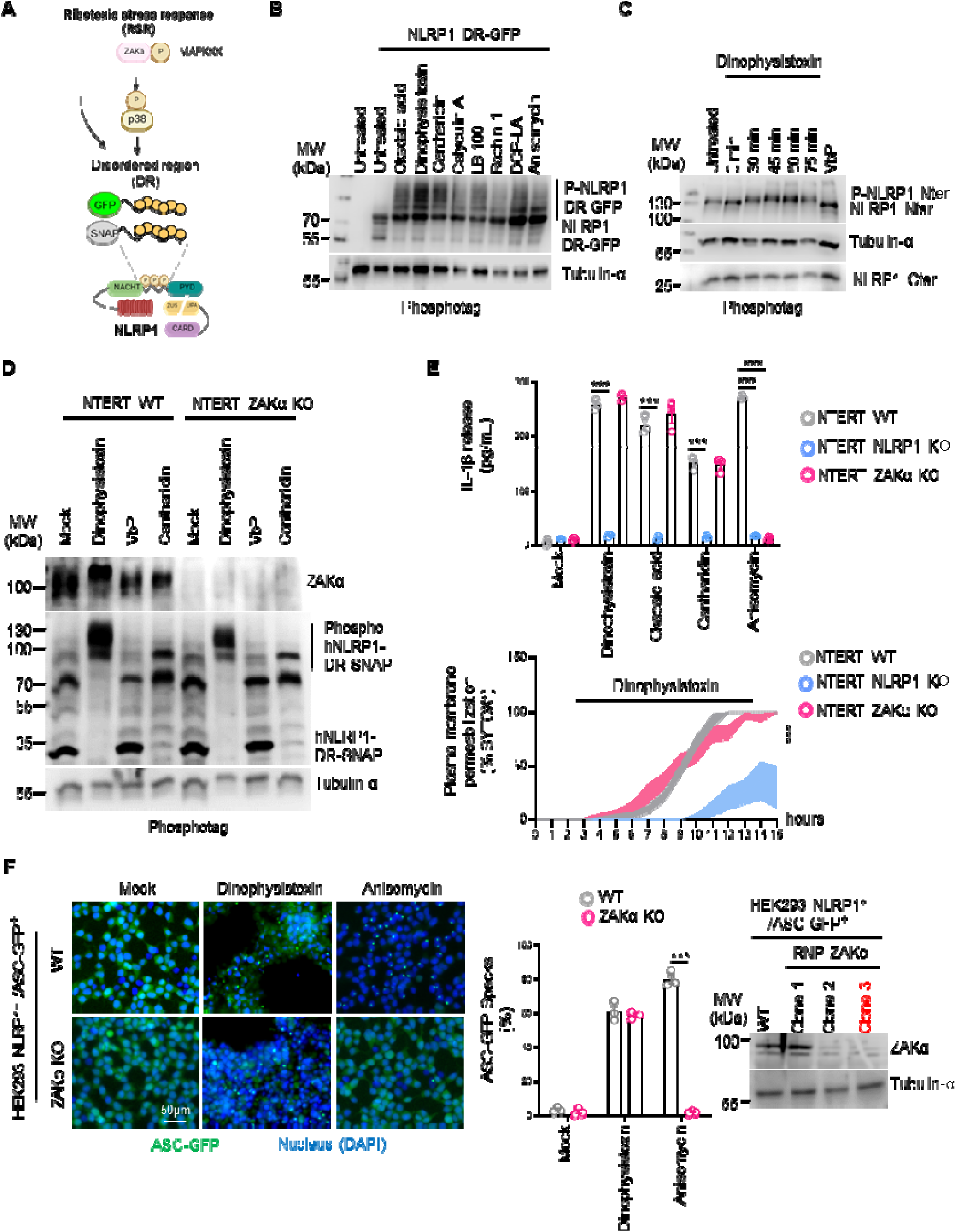
PP1/PP2 A-targeting toxins activate the NLRP1 inflammasome in a ZAKα-independent manner. **A.** Schematic representation of the mechanism of ZAKα/P38 stress kinases activation upon induction of Ribotoxic Stress Response (RSR). **B.** Phosphotag blotting of phosphorylated NLRP1 disordered Region (DR) in HEK293T expressing the NLRP1 DR construct (aa 86-275-GFP (described in **A**)) and exposed to all PP1/PP2A-targeting compounds identified in Fig. 1A**/B**) or to the known RSR inducer Anisomycin (1 µg/mL) for an hour. Tubulin-α was used as internal protein loading controls. Immunoblots show lysates from one experiment performed at least two times. **C.** Phosphotag blotting of phosphorylated full length NLRP1 in primary human keratinocytes exposed Dinophysis toxin (100 nM) for various time. Tubulin-α was used as internal protein loading controls. Immunoblots show lysates from one experiment performed at least two times. **D.** Phosphotag blotting of phosphorylated ZAKα and NLRP1 disordered Region (DR) in WT or ZAK KO NTERT NLRP1 KO + 86-275-SNAP keratinocytes exposed to Dinophysis toxin (100nM), Cantharidin (5%M) or Val-boro-Pro (VbP, 10µM) for an hour. Tubulin-α was used as internal protein loading controls. Immunoblots show lysates from one experiment performed at least three times. **E.** Plasma membrane permeabilization (SYTOX Green incorporation, 16 h) and IL-1β release evaluation (10 h) in WT, ZAKα or NLRP1 KO NTERT keratinocytes after exposure to Dinophysis toxin (100nM), Cantharidin (5%M), Okadaic acid (250nM) or Anisomycin (1µM). ***P ≤ 0.0001, one-way ANOVA. Values are expressed as mean ± SEM. Graphs show one experiment performed in triplicates at least three times. **F.** Immunoblotting and clonal selection (clone 3, red), fluorescence microscopy and associated quantifications of ASC-GFP specks in WT or ZAKα KO HEK293T^ASC-GFP/NLRP1^ reporter cells exposed to Dinophysis toxin (100nM) or Anisomycin (1µM) for 5 hours. ASC-GFP (green) pictures were directly taken in dish after adding Hoechst (nuclei staining). Images shown are from one experiment and are representative of three independent experiments; scale bars, 50 µm. ASC complex percentage was performed by determining the ratios of cells positive for ASC speckles on the total nuclei (Hoechst). At least ten fields from each experiment were analyzed. Values are expressed as mean ± SEM. ***P ≤ 0.0001, one-way ANOVA.

Phos-tag analysis revealed that all PP1/PP2A-targeting toxins and compounds triggered marked phosphorylation of hNLRP1-DR–GFP (Fig. 2B). These findings were corroborated in human TERT keratinocytes, in which full-length hNLRP1 displayed time-dependent phosphorylation following exposure to dinophysistoxin-1 (Fig. 2C). In contrast, Val-boroPro (VbP), a phosphorylation-independent activator of NLRP1, did not induce detectable hNLRP1 phosphorylation (Fig. 2C). Together, these results indicate that PP1/PP2A inhibition unleashes strong hNLRP1 phosphorylation, likely through activation of one or more yet-unidentified protein kinases.

Because the stress-activated kinase ZAKα was recently shown to drive hNLRP1 phosphorylation and activation in response to ribotoxic stress, we examined whether ZAKα contributes to NLRP1 activation induced by PP1/PP2A-targeting toxins. WT, ZAKα KO, and NLRP1 KO TERT keratinocytes expressing a SNAP-tagged hNLRP1-DR (DR-SNAP) were treated with dinophysistoxin-1, cantharidin, or VbP (Fig. 2D) (*21*). DR-SNAP phosphorylation was comparable in WT and ZAKα KO cells after toxin treatment, whereas VbP did not induce significant phosphorylation, demonstrating that toxin-induced NLRP1 phosphorylation occurs independently of ZAKα (Fig. 2D).

These results were supported by measurements of plasma membrane permeabilization (Sytox Green uptake) and IL-1β secretion in WT, ZAKα KO, and NLRP1 KO keratinocytes following PP1/PP2A inhibition (Fig. 2E). While anisomycin induced ZAKα-dependent IL-1β release, all PP1/PP2A-targeting toxins triggered IL-1β secretion and membrane permeabilization independently of ZAKα (Fig. 2E). Similarly, ASC speck formation in WT and ZAKα KO HEK293 ASC-GFP/NLRP1 cells showed that dinophysistoxin induces robust speck assembly regardless of ZAKα status (Fig. 2D).

Altogether, these results demonstrate that PP1/PP2A-targeting toxins drive strong phosphorylation of the NLRP1 disordered region, and that this process occurs independently of the stress kinase ZAKα.

### Multiple P38 MAPkinases contribute to NLRP1 inflammasome activation

Because PP1/PP2A-targeting toxins promote NLRP1 phosphorylation in a ZAKα-independent manner, we next sought to identify the kinase(s) responsible for driving the hNLRP1 inflammasome response. To this end, we performed a PAMGENE kinase activity assay in human keratinocytes exposed to PP1/PP2A-targeting toxins (Fig. 3A). This analysis revealed robust activation of multiple MAP kinase family members (p38α, p38β, p38δ, JNK1/2, ERK1/2), NF-κB–related kinases (TBK1, IKKε), as well as CAM kinases and several cell-cycle kinases (Fig. 3A).

**Figure 3.**
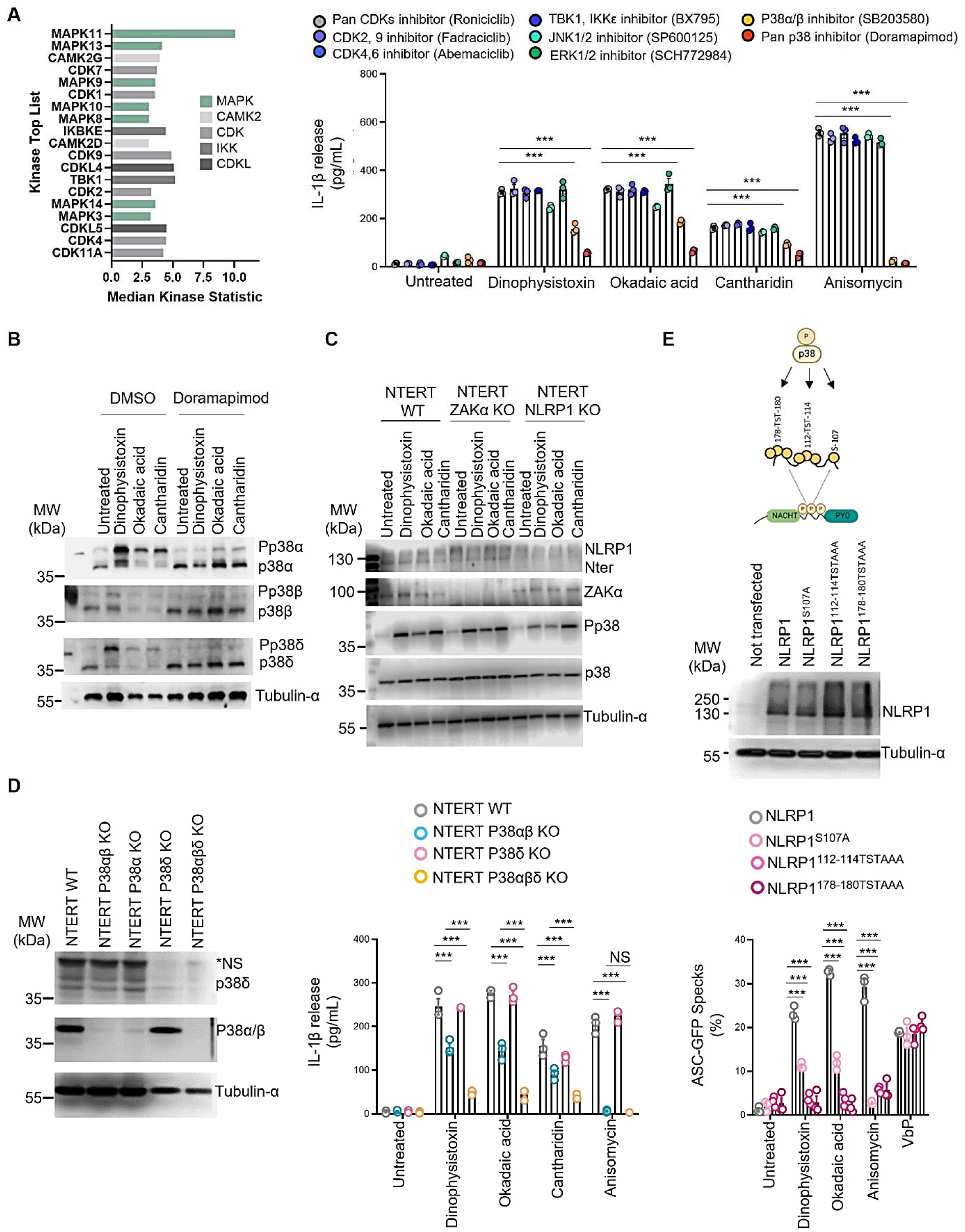
Multiple P38 MAPkinases contribute to NLRP1 inflammasome activation. **A.** Pamgene analysis of activated Serine/threonine kinases in primary human keratinocytes exposed to Dinophysis toxin (100nM) for 2 hours and subsequent determination of IL-1β release in WT NTERT-keratinocytes after 8 h exposure to Dinophysis toxin (Dino. toxin, 100nM), Okadaic acid (Ok. Acid, 250nM), Cantharidin (Canth. 5µM) or Anisomycin (1µM) in presence/absence of inhibitors of identified kinases in. For all kinases, inhibitors were used at 10µM. ***P ≤ 0.0001, one-way ANOVA. Values are expressed as mean ± SEM. Graphs show one experiment performed in triplicates at least two times. **B.** Phosphotag blotting of phosphorylated P38 kinase isoforms in NTERT keratinocytes exposed to Dinophysis toxin (Dino. toxin, 100nM), Okadaic acid (Ok. Acid, 250nM), Cantharidin (Canth. 5µM) for 1 hour in presence/absence of the Pan P38 inhibitor Doramapimod (10µM). Tubulin-α was used as internal protein loading controls. Immunoblots show lysates from one experiment performed at least three times. **C.** Immunoblotting of P38, ZAKα, NLRP1 and phosphorylated P38 kinases in WT, ZAKα KO or NLRP1 KO NTERT keratinocytes exposed to Dinophysis toxin (Dino. toxin, 100nM), Okadaic acid (Ok. Acid, 250nM), Cantharidin (Canth. 5µM) for 1 hour. Tubulin-α was used as internal protein loading controls. Immunoblots show lysates from one experiment performed at least three times. **D.** Immunoblotting characterization of the P38 isoform genetic knockdown (CRISPR-Cas9) and of the subsequent IL-1β release in WT, P38δ KO, P38α/β dKO, or P38α/β/δ TKO NTERT keratinocytes exposed or not to Dinophysis toxin (Dino. toxin, 100nM), Okadaic acid (Ok. Acid, 250nM), Cantharidin (Canth. 5µM) or Anisomycin (1µM) for 8 hours. ***P ≤ 0.0001, one-way ANOVA. Values are expressed as mean ± SEM. Graphs show one experiment performed in triplicates at least three times. **E.** Western blot showing NLRP1 (anti-NLRP1 N-terminal antibody (aa 1–323)) and associated fluorescence microscopy/quantifications of ASC-GFP specks in HEK293^ASC-GFP^ reporter cells reconstituted with hNLRP1 or hNLRP1 plasmid constructs mutated for important 38 phosphorylation sites (S107A, TST112-114AAA and TST178-180AAA) after 10 h of exposure to Dinophysis toxin (100nM), Okadaic acid (250nM), Cantharidin (5µM) or Val-boro-Pro (VbP, 10µM). ASC-GFP (green) pictures were taken in the dish after toxin exposure. Images shown are from one experiment and are representative of three independent experiments; scale bars, 10 µm. ASC complex percentage was performed by determining the ratios of cells positive for ASC speckles (green, GFP) on the total nuclei (Hoechst). At least ten fields from three independent experiments were analyzed. Values are expressed as mean ± SEM. ***P ≤ 0.0001, two-way ANOVA with multiple comparisons. Graphs show one experiment performed in triplicate at least three times.

We next assessed the contribution of these kinase families to hNLRP1 regulation using a pharmacological approach (Fig. 3B). Inhibitor screening showed that the p38α/β inhibitor SB203580 and the pan-p38 inhibitor doramapimod (*30*) both reduced IL-1β release in TERT keratinocytes treated with PP1/PP2A-targeting toxins, suggesting that p38 kinases are involved in NLRP1 inflammasome activation (Fig. 3B). In contrast, inhibition of the other activated kinases did not significantly affect toxin-induced IL-1β release. The greater efficacy of doramapimod compared with SB203580 further suggested that additional p38 isoforms beyond p38α/β contribute to this process.

Consistent with this idea, our PAMGENE data also identified p38δ as strongly activated in keratinocytes exposed to dinophysistoxin (Fig. 3A). Phos-tag analyses confirmed that all three p38 isoforms, p38α, p38β, and p38δ, undergo robust phosphorylation in response to PP1/PP2A-targeting toxins (Fig. 3B, C). These results prompted us to assess the individual and collective roles of these isoforms in hNLRP1 activation. Using CRISPR-Cas9, we generated single-isoform knockout TERT keratinocytes and measured IL-1β release following PP1/PP2A inhibition (Fig. 3D). Loss of p38δ or p38α/β alone caused minimal or only modest reductions in IL-1β release, whereas combined deletion of all three isoforms (p38 TKO) led to a strong, though incomplete, inhibition (Fig. 3D). These findings indicate that p38α, β, and δ cooperatively regulate hNLRP1 inflammasome activation in response to PP1/PP2A-targeting toxins.

To further validate the requirement for p38 kinases in this pathway, we used reporter cell lines expressing either WT hNLRP1 or p38-insensitive hNLRP1 variants in which important p38-responsive residues in the disordered region (Ser107 and Thr-Ser-Thr 112–114/178–180) were replaced with alanines (A) (Fig. 3E) (*20*, *22*). ASC speck formation assays showed that PP1/PP2A inhibition failed to induce speck assembly in cells expressing p38-insensitive hNLRP1, whereas cells expressing WT hNLRP1 displayed robust ASC oligomerization (Fig. 3E). Importantly, Val-boroPro (VbP), a p38-independent activator of NLRP1, induced similar ASC speck formation in all cell lines, confirming that the introduced mutations do not impair other activation pathways (Fig. 3E).

Together, these results demonstrate that the p38α, β, and δ isoforms act collectively as key regulators of hNLRP1 inflammasome activation following PP1/PP2A inhibition.

### TAK1 apical MAP kinase both activates P38 kinases and directly contributes to triggering the hNLRP1 inflammasome in various contexts

Our observation that multiple p38 isoforms (p38α, β, and δ) contribute to hNLRP1 activation in response to PP1/PP2A-targeting toxins led us to hypothesize that PP1/PP2A might regulate upstream MAP kinases involved in p38 activation. To test this, we examined whether any apical MAP3Ks participate in the hNLRP1 inflammasome response induced by PP1/PP2A inhibition. We performed a pharmacological screen targeting the major druggable MAP3Ks in the HEK293 ASC-GFP/NLRP1 reporter system (Fig. 4A). Strikingly, only HS-276, a preclinical inhibitor of the MAP3K7 kinase TAK1 (*31*), robustly suppressed ASC speck formation following toxin exposure (Fig. 4A), suggesting that TAK1 is a key upstream regulator of hNLRP1 activation under PP1/PP2A-inhibiting conditions.

**Figure 4.**
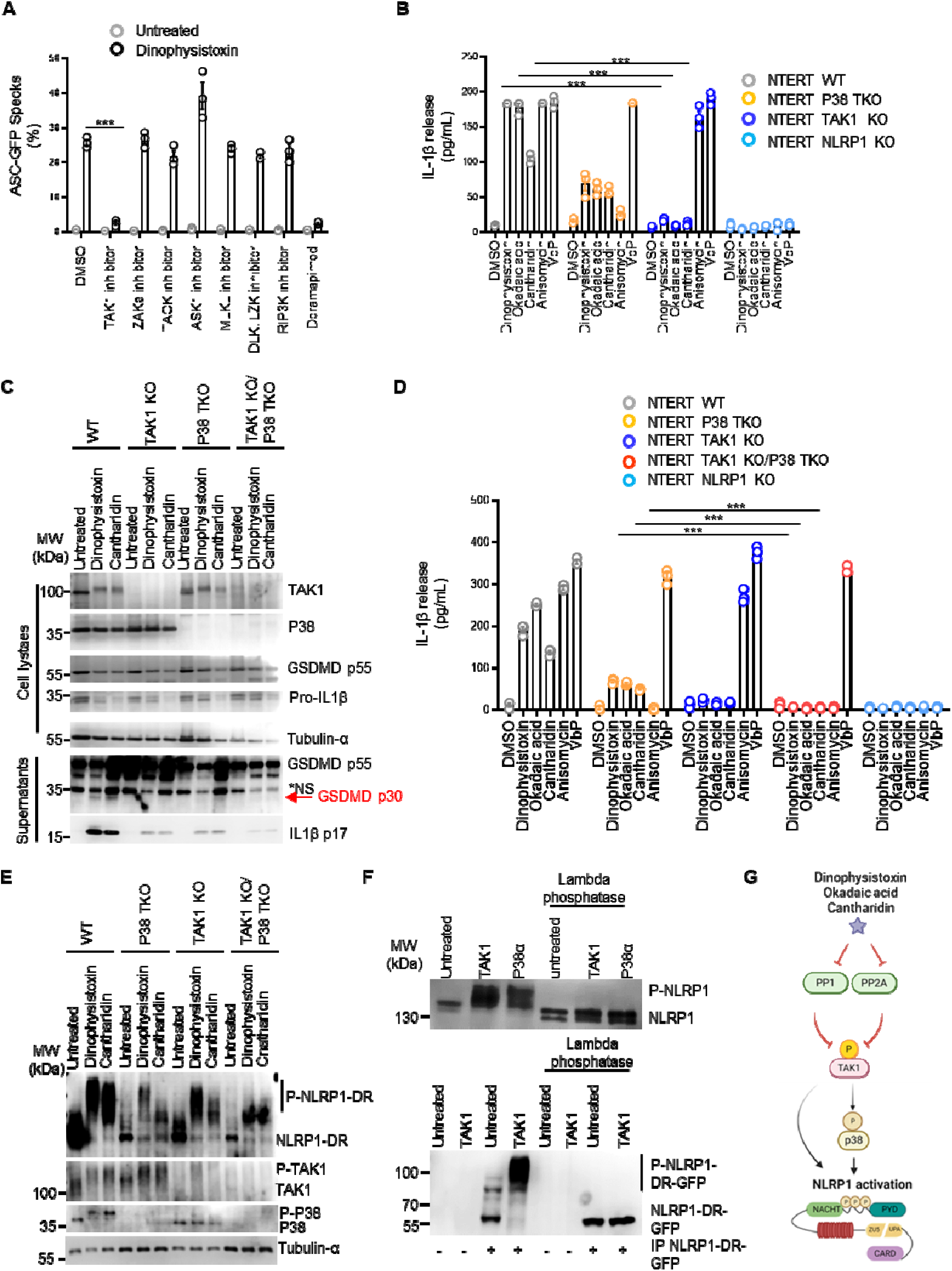
TAK1 apical MAP kinase both activates P38 kinases and directly contributes to triggering of the hNLRP1 inflammasome. **A.** Quantifications of ASC-GFP specks in HEK293^ASC-GFP/NLRP1^ reporter cells exposed to Dinophysis toxin (100nM) or not for 6 h in presence or absence of various MAPK inhibitors (10µM). TAK1 inhibitor; HS-276, ZAKα inhibitor; PLX4720, TAOK inhibitor; CP-43, MLKL inhibitor; necro sulfonamide, ASK1 inhibitor; GS-444217, DLK/LZK inhibitor; DN-1289, RIPK3 inhibitor; GSK-872. ASC-GFP (green) pictures were directly taken in dish after adding Hoechst (nuclei staining). Images shown are from one experiment and are representative of three independent experiments; scale bars, 10 µm. ASC complex percentage was performed by determining the ratios of cells positive for ASC speckles on the total nuclei (Hoechst). At least ten fields from each experiment were analyzed. Values are expressed as mean ± SEM. ***P ≤ 0.0001, one-way ANOVA. **B.** Determination of the IL-1β release in WT, P38α/β/δ TKO, TAK1 KO and NLRP1 KO NTERT keratinocytes exposed or not to Dinophysis toxin (Dino. toxin, 100nM), Okadaic acid (Ok. Acid, 250nM), Cantharidin (Canth. 5µM), Anisomycin (1µM) and VbP (10µM) for 10 hours. ***P ≤ 0.0001, one-way ANOVA. Values are expressed as mean ± SEM. Graphs show one experiment performed in triplicates at least three times. **C.** Immunoblotting of P38, TAK1, cleaved GSDMD and IL-1β and of the subsequent IL-1β release in WT, P38α/β/δ TKO, TAK1 KO, P38α/β/δ TKO/TAK1 KO and NLRP1 KO NTERT keratinocytes exposed or not to Dinophysis toxin (Dino. toxin, 100nM), Okadaic acid (Ok. Acid, 250nM), Cantharidin (Canth. 5µM), Anisomycin (1µM) and VbP (10µM) for 10 hours. ***P ≤ 0.0001, one-way ANOVA. Values are expressed as mean ± SEM. Graphs show one experiment performed in triplicates at least three times. **D.** Phosphotag blotting of phosphorylated P38, TAK1 and NLRP1-DR-SNAP in WT, P38α/β/δ TKO, TAK1 KO or P38α/β/δ TKO/TAK1 KO NTERT keratinocytes exposed to Dinophysis toxin (Dino. toxin, 100nM), Okadaic acid (Ok. Acid, 250nM), Cantharidin (Canth. 5µM) for 1 hour. Tubulin-α was used as internal protein loading controls. Immunoblots show lysates from one experiment performed at least three times. **E.** Phosphotag immunoblotting of phosphorylated recombinant NLRP1 full-length protein or immunoprecipitated GFP-tagged NLRP1-DR incubated with recombinant TAK1-TAB1 fusion or P38α kinases for 60 minutes in presence/absence of lambda phosphatase. Immunoblots show proteins from one experiment performed at least three times. **F.** Fluorescence microscopy quantifications of ASC-GFP specks in WT, P38α/β/δ TKO or in P38α/β/δ TKO/TAK1 KO HEK293^ASC-GFP/NLRP1^ reporter cells after 6 h of exposure to Dinophysis toxin (100nM) or Val-boro-Pro (VbP, 10µM). ASC complex percentage was performed by determining the ratios of cells positive for ASC speckles (green, GFP) on the total nuclei (Hoechst). At least ten fields from three independent experiments were analyzed. Values are expressed as mean ± SEM. ***P ≤ 0.0001, two-way ANOVA with multiple comparisons. Graphs show one experiment performed in triplicate at least three times.

To validate this genetically, we disrupted TAK1 in TERT keratinocytes using CRISPR–Cas9 and assessed IL-1β release after stimulation with dinophysistoxin, okadaic acid, or cantharidin (Fig. 4B). While anisomycin and VbP elicited strong IL-1β secretion in both wild-type and TAK1-deficient cells, thus confirming their TAK1-independent mode of action, TAK1 KO cells failed to release IL-1β in response to any PP1/PP2A-targeting toxin (Fig. 4B), establishing TAK1 as a central mediator of toxin-induced NLRP1 inflammasome activation.

Because TAK1 is also known to activate p38 kinases (*32*, *33*), we next asked (1) whether PP1/PP2A inhibition leads to TAK1 activation and (2) whether TAK1 is required for toxin-induced p38 activation (Fig. 4C–E). We found that all three toxins induced robust TAK1 phosphorylation (Fig. 4C–E). Moreover, toxin-induced p38 phosphorylation was completely abolished in TAK1 KO cells (Fig. 4E), indicating that TAK1 not only becomes activated following PP1/PP2A inhibition but also is essential for downstream p38 activation.

To dissect the respective roles of TAK1 and p38 in NLRP1 inflammasome activation, we evaluated IL-1β release, NLRP1-DR phosphorylation, and gasdermin-D cleavage (Fig. 4C–E). Both p38 TKO and TAK1 KO keratinocytes showed markedly reduced IL-1β production and gasdermin-D processing in response to the toxins (Fig. 4C, D). Notably, the reduced IL-1β production was stronger in TAK1 KO than in p38 TKO cells, indicating that TAK1 exerts both p38-dependent and p38-independent effects on hNLRP1 activation (Fig. 4C, D). Consistently, combined deletion of TAK1 in the p38 TKO background (p38 TKO/TAK1 KO) resulted in a complete loss of toxin-induced IL-1β release and gasdermin-D cleavage (Fig. 4C, D). In addition, NLRP1-DR phosphorylation was slightly impaired in either TAK1 KO or p38 TKO cells, and further reduced in p38 TKO/TAK1-KO cells (Fig. 4E), supporting a cooperative yet partially independent contribution of p38 and TAK1 kinases.

Given previous work showing that the apical kinase ZAKα can directly phosphorylate NLRP1, we next asked whether TAK1 might also directly phosphorylate hNLRP1 in addition to activating p38. Using recombinant active TAK1 and p38α, we found that both kinases phosphorylated full-length hNLRP1 and its DR region *in vitro* (Fig. 4F), suggesting that TAK1 can directly phosphorylate and activate the hNLRP1 inflammasome (Fig. 4G).

The strong involvement of TAK1 in toxin-induced hNLRP1 activation led us to examine its potential role in other NLRP1-activating contexts. Alphavirus infections and dsRNA have been shown to induce a phosphorylation-dependent, albeit only partially ZAKα-dependent, NLRP1 response, suggesting the involvement of additional apical kinases (*19*, *20*). We therefore hypothesized that TAK1 could contribute to dsRNA- and virus-induced NLRP1 activation in keratinocytes. Wild-type, TAK1 KO, ZAKα KO, p38 TKO, p38 TKO/TAK1 KO, and NLRP1 KO TERT keratinocytes were either transfected with high-molecular-weight dsRNA (poly:IC) or infected with VSV, oncolytic VSV-M51, or Sindbis virus (Fig. S4A, B) (*20*). At the exception of VSV, all stimuli triggered NLRP1-dependent IL-1β release (Fig. S4A, B). Whereas single loss of ZAKα, TAK1, or of the p38 isoforms partially reduced IL-1β secretion, combined TAK1 KO/p38 TKO or ZAKα KO/TAK1 KO cells completely impaired IL-1β release upon dsRNA transfection or viral infection (Fig. S4A, B). Thus, these results suggest that TAK1 and TAK1-associated pathways are key additional regulators of the hNLRP1 inflammasome in keratinocytes.

Because PP1/PP2A inhibition activates TAK1, we finally tested whether PP2A activation might conversely dampen TAK1-mediated NLRP1 signaling. Using the PP2A activator ATUX-1215 in ZAKα KO A549 ASC-GFP/NLRP1 reporter cells transfected with dsRNA, we observed that PP2A activation strongly reduced ASC speck formation during dsRNA stimulation (Fig. S4C). To the contrary, PP2A activation had no effect on VbP-induced NLRP1 activation (Fig. S4C). Hence, these results suggest that PP2A activity might specifically increase the activation threshold of the TAK1-dependent NLRP1 pathway upon dsRNA exposure.

Altogether, our findings show that PP1/PP2A-targeting toxins and dsRNA converge on TAK1 activation, which in turn drives both p38-dependent and p38-independent activation of the hNLRP1 inflammasome in keratinocytes.

### TAK1 and P38 kinases play a major role in hNLRP1 inflammasome response in Human native skin models of toxin exposure

The identification of TAK1 as a major contributor to PP1/PP2A-targeting toxin–induced hNLRP1 inflammasome activation prompted us to examine the relevance of this pathway in a more physiologically complex setting. To this end, we established a model of human native skin explants exposed to the dermatitis-inducing toxins cantharidin, dinophysistoxin, and okadaic acid (*34*). These explants, obtained from healthy donors, contain the full pattern of skin-resident cell types including keratinocytes, fibroblasts, melanocytes, mast cells, and Langerhans cells, thus providing a biologically representative environment (*34*).

In this model, all three toxins induced pronounced intraepidermal adhesion loss and extensive cell death (Fig. 5A). Notably, pretreatment with the TAK1 inhibitor HS-276 or the pan-p38 inhibitor doramapimod markedly preserved intradermal adhesion and overall tissue architecture in explants exposed to dinophysistoxin or okadaic acid (Fig. 5A-C). This finding suggests that both TAK1 and p38 kinases are key mediators of PP1/PP2A-targeting toxin–induced skin damage. In contrast, the protective effect of these inhibitors against cantharidin-induced injury was less pronounced (Fig. 5A), suggesting that cantharidin may elicit skin pathology through both PP1/PP2A-dependent and -independent mechanisms (*35*).

**Figure 5.**
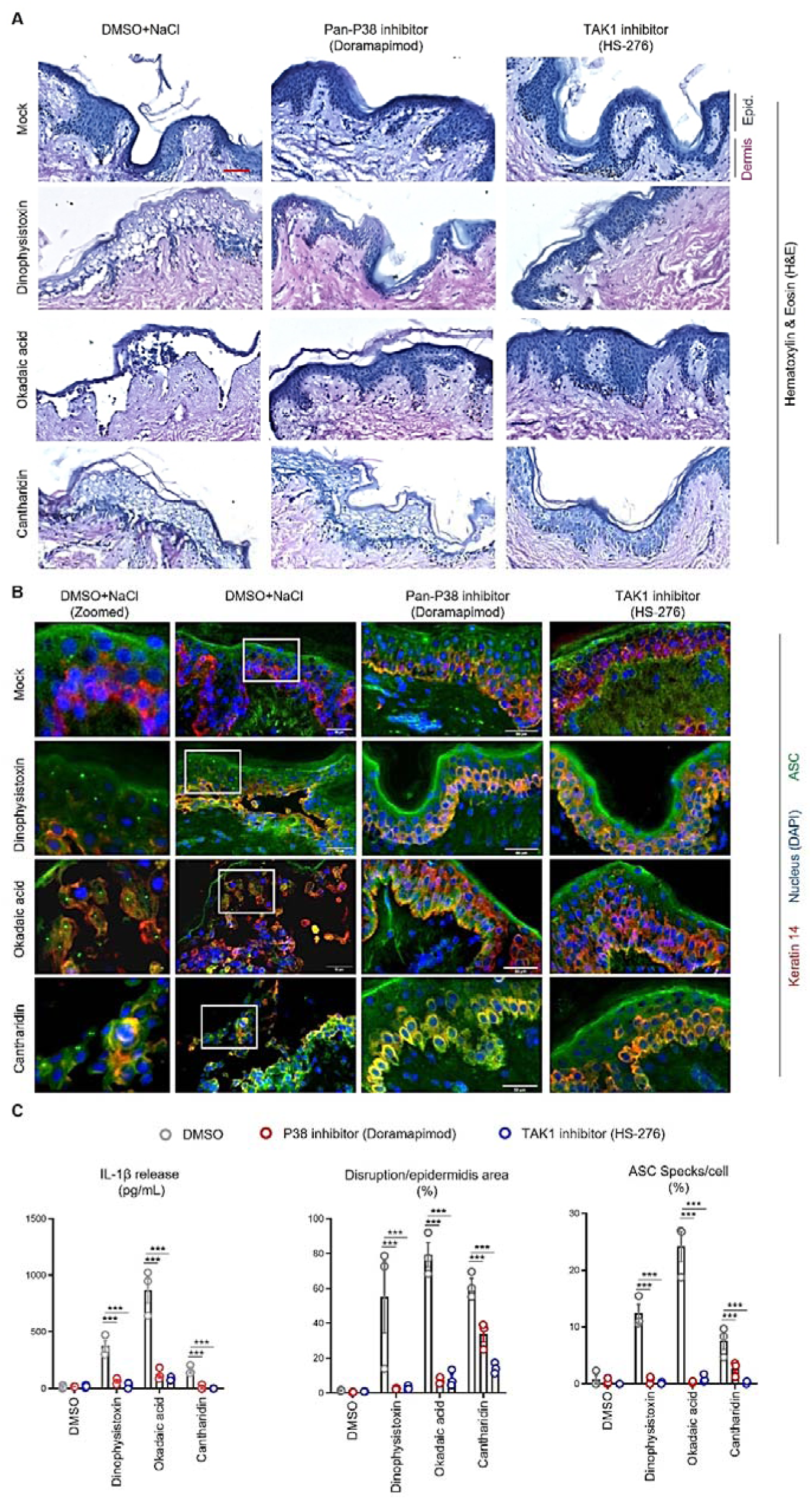
TAK1 and P38 kinases play a major role in hNLRP1 inflammasome response in Human native skin models of toxin exposure. **A-C**. Hematoxylin (H) & Eosin (E) or ASC immunobiological staining showing P38 and TAK1-dependent histological/inflammasome changes caused by Dinophysis toxin (250 nM), Okadaic acid (600 nM) and Cantharidin (10µM) exposure for 24 hours. When specified, pan P38 kinase inhibitor Doramapimod (10µM) and TAK1 inhibitor HS-276 (20µM) were used. Associated quantification of IL-1β release, the dermal–epidermal layer detachment and the percentage of ASC specks in the Human skin explants. P values indicated in figure, one-way ANOVA. Images are representative of two (A) and three (B) biological replicates. Scale bar = 50 µm.

We next assessed whether these toxins could trigger an inflammasome response within the explants. Longitudinal analysis of IL-1β release revealed robust cytokine production beginning 24 hours after exposure, consistent with an inflammasome activation (Fig. S5A). Immunohistochemistry further demonstrated abundant ASC specks in keratin 14-positive keratinocytes (Fig. 5B), and proximity ligation assays showed close spatial association between ASC and NLRP1 (Fig. S5B). Together, these data suggest that cantharidin, dinophysistoxin, and okadaic acid all activate the NLRP1 inflammasome in human skin. Finally, to define the contribution of TAK1 and p38 kinases to this response, we pretreated skin explants with HS-276 or doramapimod prior to toxin exposure. Both inhibitors markedly reduced IL-1β release and ASC speck formation across all three toxins (Fig. 5B, C).

All in one, these results demonstrate that TAK1 and p38 kinases are not only critical for NLRP1 inflammasome activation in response to PP1/PP2A-inhibiting toxins but also for the associated tissue damage.

## DISCUSSION

Because of its strong sensitivity to phosphorylation-driven activation, we investigated the ability of cellular phosphatases to modulate the NLRP1 inflammasome response. By performing a phosphatase screening approach, we identified PP1 and PP2A phosphatases as critical inhibitors of the NLRP1 inflammasome response. Specifically, genetic inactivation of PP1/PP2A catalytic subunits or exposure to PP1/PP2A-inhibiting toxins that can cause skin dermatitis, such as Cantharidin from the beetles’ blister or Okadaic acid and Dinophysis toxins from *Dinophysis* algae, all triggered the MAP3K7 TAK1, which promoted a phosphorylation cascade that culminated with the NLRP1 inflammasome activation in human keratinocytes. Given their presence in various living organisms such as viruses (e.g. small antigen T) (*36*, *37*), insects (*29*), algal species (*26*), PP1/PP2A-inhibiting toxins may represent a broad, yet poorly characterized, family of triggers of NLRP1 activation with potential dermatologic, gastrointestinal, or respiratory manifestations. However, in a coevolution point of view, it appears unlikely that the NLRP1 inflammasome has been selected by the existence of various PP1/PP2A-targeting toxins, given their presence in very distinct and specific geographical areas (*26*, *27*, *29*). To this regard, our findings along the ones from two companion studies from Chua et *al* and Corley et *al*., found that TAK1 kinase also contributes to the NLRP1 inflammasome response upon double stranded RNA (dsRNA) exposure or viral infections driven by the oncolytic virus VSV-M51 and Sindbis viruses. Of importance, although toxin-inactivated PP1/PP2A solely promotes a TAK1-dependent NLRP1 response, dsRNA and viral infections engage multiple activating pathways including but not restricted to 1/ NLRP1 binding, 2/ Ribotoxic stress response-driven ZAKα activation and 3/ TAK1 hyperactivation, suggesting that yet to be discovered and characterized effectors/signaling pathway might be at play in cells exposed to viruses/dsRNA (*19*, *20*).

Our findings that PP1 and PP2A phosphatase regulate the human NLRP1 inflammasome through TAK1 also underscore the versatility of NLRP1 as a sensor of diverse threats. In this context, an emerging picture suggests that NLRP1 is positioned as a central integrator of stress signals originating from infectious agents, environmental toxins, and chemical disruptors that all converge on MAPK signaling and regulation (*38*). To this regard, we hypothesize that PP1/PP2A but also additional phosphatases could be acting as counterbalances against multiple MAPK activity under homeostatic or stress conditions (*39*), hence increasing the NLRP1 inflammasome activation threshold, as observed in our study during viral infections and dsRNA exposure. An additional hypothesis of work is that phosphatases of the host, including PP1/PP2A, might also be involved at directly dephosphorylating NLRP1 disordered region (DR) in addition to inactivating kinases, hence fostering the cells to avoid NLRP1 spontaneous activation. Although we did not address specifically this question in our study, this is currently under investigations in our group. This is of particular relevance for understanding how keratinocytes, the primary cells of the epidermis, maintain immune homeostasis while being exposed to a constant barrage of environmental toxins and pathogens.

While our study provides significant insights into the role of PP1 and PP2A phosphatases in the regulation of NLRP1 inflammasome activation in keratinocytes, several important questions remain unanswered, and further research is necessary to fully understand the mechanistic details and broader implications of these findings. To this regard, although we demonstrated the critical involvement of several of the catalytic subunits of PP1 and PP2A in regulating NLRP1 inflammasome activation, the specific adaptor proteins and subunits that mediate this effect remain unclear. Indeed, PP1 and PP2A consist of multiple isoforms with distinct tissue-specific functions associated with a multitude of different adaptor proteins that confer to those holoenzymes their specificity of action (*23–25*, *37*). Future studies should focus on characterizing the individual subunits of PP1 and PP2A that mediate their inhibitory function on TAK1-mediated NLRP1 inflammasome regulation, as well as the potential role of regulatory subunits in fine-tuning inflammasome dephosphorylation.

Furthermore, the strong importance of PP1/PP2A at restricting kinase-driven NLRP1 phosphorylation and activation pushes us to speculate that multiple viruses, bacteria, fungi or parasites might have set up phosphatase or phosphatase-like virulence factors that in theory could dampen the NLRP1 inflammasome response (*40*), either by targeting key MAPK and/or by directly dephosphorylating NLRP1 (*41*), which will constitute an exciting future area of research.

Our study further delves into the kinase networks that drive NLRP1 inflammasome activation following PP1/PP2A inhibition. Specifically, we found that TAK1 could mediate both P38 kinase-dependent and P38-independent activation of the NLRP1 inflammasome. To this regard, it is worth mentioning that among the P38 kinases activated by PP1/PP2A-targeting toxins were the multiple P38 isoforms (α, β, and δ), which suggests a strong redundancy among those kinases at promoting NLRP1 phosphorylation and activation. Those findings join the recent study from Jenster et al., that also demonstrated a key role of all P38 kinase isoforms on NLRP1 phosphorylation and activation (*20*). In line with this, Chua et al. (companion manuscript) found a minor role for P38α/β isoforms in Cantharidin-induced NLRP1 inflammasome response. We hypothesize that, given the redundant impact of the three P38 isoforms on the NLRP1 inflammasome response, P38α/β KO might not be sufficient to measure the contribution of P38 or that under certain conditions TAK1 might overwhelm P38 kinase requirement.

Regarding the relevance of our findings in complex settings, our Human native skin explant approaches highlighted the strong contribution of both TAK1 and P38 kinases on the NLRP1 inflammasome activation in response to PP1/PP2A-targeting toxins. Importantly, pretreatment with TAK1 or P38 inhibitors also significantly reduced skin damage, thus suggesting that these kinases contribute not only to inflammasome signaling but also to the inflammatory skin pathology observed in response to toxin exposure. Whether additional physiological or pathological contexts involve dysregulated P38/TAK1 kinases such as psoriasis, hypomorphic PP1/PP2A conditions (*42*) or genetic deficiency in RNA regulation (e.g. ADAR1 alterations) (*43*) will warrant for further studies and testing preclinical P38/TAK1 inhibitors in a topical way. To the contrary, such formulation of PP1/PP2A inhibition, when added locally holds strong potential to act as a curing agent. To this regard, Cantharidin is the main compound of the YCanth cure, commonly used to eliminate local warts induced by the molluscum virus in the skin (*44*). It is tempting to speculate that such process might involve NLRP1-driven pyroptosis and inflammation in treated cells.

### Limitations of the study

Although our study identifies PP1/PP2A phosphatases as critical regulators of the NLRP1 inflammasome, we did not address the involvement of the multiple adaptors that drive the specificity of those holoenzymes, which is currently under extensive study in the lab. Furthermore, despite the use of human native skin explants from healthy donors, our study lacks the complete physiology of an organism. Given the lack of conservation between rodents and humans on the NLRP1 phosphorylation, such approach is at this time not available as we have not yet succeeded at generating robust humanized models.

## MATERIALS AND METHODS

### Study design

This study aimed to define how phosphorylation controls human NLRP1 inflammasome activation. We first screened phosphatases inhibitors and environmental phosphatase-targeting toxins in reporter cells. Anisomycin and Val-boro-Pro, two known activators of NLRP1 were used as a treatment of reference. The involvement of the Ribotoxic Stress Response and ZAKα was examined in A549, HEK293T and N/TERT keratinocytes through pharmacological inhibition and CRISPR-Cas9 depletion. Candidates’ kinases were investigated by kinase activity assay, inhibitor profiling, genetic knock-out, and *in-vitro* kinase and lambda protein phosphatase assays coupled to PhosTag SDS-PAGE. The requirement for TAK1 upstream kinase in NLRP1 activation by dsRNA and viral infection was evaluated in keratinocytes stimulated with poly(I:C) or infected with various viral strains, including VSV or VSV-M51, or Sindbis virus. Finally, toxin-induced NLRP1 activation and kinase dependency were validated in human skin explants using pharmacological inhibitors and histology, immunofluorescence, proximity ligation assays and cytokine measurements. Details on sample size and statistical analyses are provided in figure legends.

### Human Skin Tissue Models

HypoSkin 22-mm-diameter human skin explants (GenoSkin, Toulouse, France) derived from abdominal tissue were obtained from two healthy adult Caucasian donors (female, 35-year-old; male, 43-year-old), both of whom had no documented dermatological conditions and gave written informed consent. This biological material complies fully with the Declaration of Helsinki and all relevant regulations. The study is not classified as human subject research, and Institutional Review Board approval was not required. Each donor batch underwent the manufacturer’s internal quality control prior to shipment, including histological validation of tissue integrity and virological screening for HIV-1 and −2 and Hepatitis B and C.

Explants were maintained in the recommended culture medium (GenoSkin, NSMED2) and standard culture conditions (37°C, 5% CO_2_). Culture medium was replaced daily. When indicated, inhibitors (Doramapimod, 20µM, MedChem Express, HY-10320; TAK-1 inhibitor HS-276 (20 µM, MedChem Express, HY-147141) were added to fresh medium and applied as a 24-hour pre-treatment before injection. Each explant received a 55µL intradermal injection delivered just beneath the epidermis. Injection solutions contained either phosphatase-targeting toxins (Dinophysistoxin, 500nM, Bertin Technologies, WP-10011497; Okadaic acid, 2400nM, Bertin Technologies, WP-10011490; Cantharidin, 25µM, Sigma-Aldrich, C7632-25MG) diluted in PBS, PBS, or toxin/inhibitor mixtures. Explants were collected 72 hours later, fixed in paraformaldehyde (PFA) for histological staining or snap-frozen in liquid nitrogen for immunohistochemical analyses. All experiments were performed in two independent runs, each including three biological replicates per condition.

### Cell culture

The complete description of all cells and cell lines used in this study is provided in **Table S1**. Human Embyonic Kidney (HEK) 293T and human alveolar basal epithelial (A549) cells were cultured in Dulbecco’s Eagme’s Medium (DMEM, high-glucose, Gibco, 41965039) supplemented with 10% Fetal Bovine Serum (FBS, Cytiva, SV30160.03IR).

Immortalised human keratinocytes (N/TERT) were provided by J. Rheinwald (Material Transfer Agreements to FL. Zhong and E. Meunier). WT, ZAKα KO and NLRP1 KO cells were previously described by (*21*, *22*) and WT cells stably expressing NLRP1 Disordered Region (DR) construct (aa 86-275) expressing a SNAP tag (here referred to as N/TERT + 86-275-SNAP) were generated in a previous study (*21*, *22*). N/TERT cells were cultivated in Keratinocyte Serum Free Media (SFM, Gibco, 17005042) supplemented with final concentration of 25mg/L Bovine Pituitary Extract (Gibco, 13028-014), 294.4 ng/L human recombinant EGF (Gibco, 10450-013) and 300µM of CaCl2 (PromoCell, C-34006). Human primary keratinocytes (pHEKs, PromoCell, C-12007) were cultivated in Keratinocyte Growth Medium 2 (PromoCell, C-20011).

N/TERT keratinocytes, HEK293T cells, A549 cells and pHEKs were cultured under standard conditions, at 37°C, 5% CO_2_.

### CRISPR/Cas 9 KO generation with electroporation

CRISPR/Cas9 ribonucleoprotein (RNP) complexes were prepared by mixing recombinant Cas9 protein (Invitrogen, A36498) with custom-designed sgRNA (benchling.com, SYNTHEGO) at a 1:1 molar ratio in Genome Editing Buffer (Invitrogen, A5430001). Complexes were incubated at room temperature for 25 minutes. Cell pellets were resuspended in Genome Editing Buffer (7µL buffer per 1.5 × 10^5^ cells), supplemented with GFP mRNA (1µg, OZ Biosciences, MRNA15-100) to assess electroporation efficiency, and combined with RNPs. Electroporation was performed according to the manufacturer’s protocol using 1700 V, 20 ms, single-pulse setting. Cells were immediately transferred into pre-warmed (37°C) 12-well plates and cultured for 48 hours before selection. Knock-out efficiency was confirmed by immunoblotting and further experiments were performed after one week of recovery.

The following sites were targeted: MAPK14 (p38α) (5′-TTGTGTCAAAAGCAGCACTA-3′ / 5′-AAAGAACCTACAGAGAACTG-3′); MAPK11 (p38β) (5′-GCAGAACGTACCGGGAGCTG-3′ / 5′-CCGGGCGTCGTAGGCCGAAC-3′); MAPK13 (p38δ) (5′-CTCCCCTGACCGCTTGTCGA-3′ / 5′-CAGCAGCAGCAGCTCCCGGT-3′); PPP1CA (5′-GGCTCACCGCAGATCTTGAG-3′ / 5′-GCGCCCCAGTGCAGGGCTCG-3′); PPP1CB (5′-ATTTATCGTTTGTCAGTACG-3′ / 5′-TACATACCACAAATTTTCAG-3′); PPP1CC (5′-GAAGCACCACTCAAAATATG-3′ / 5′-ATTTTTCTCTTGCAGTGAGA-3′); PPP2CA (5′-ATAAACAAGTAATTTGTATC-3′ / 5′-ACATCGAACCTCTTGCACGT-3′); PPP2CB (5′-GGATACAAACTACTTATTCA-3′ / 5′-AAAAGAATCAAATGTGCAAG-3′).

### Plasmid and nucleic acid transfection

For plasmid expression, DNA (1 µg/µL) were diluted 1:10 in nuclease-free water and mixed with LyoVec (1:100 ratio, InvivoGen, lyec-1). The DNA-LyoVec solution was gently mixed by pipetting and incubated at room temperature for 20 minutes. The transfection mixture was added dropwise to cells in antibiotic-free medium, and incubated at 37°C, 5% CO_2_. In all, 10 ng of previously described NLRP1 plasmid (pSelect2B-hNLRP1 WT, pSelect2B-hNLRP1 Ser107A, pSelect2B-hNLRP1 ^112^TST^114^/^112^AAA^114^Ser107Pro, pSelect2B-hNLRP1 ^178^TST^180^/^178^AAA^180^) (*18*, *21*) were transfected per well. Cells were analysed 48 hours after transfection.

For Poly(I:C) (Invivogen, tlrl-picw) stimulation, keratinocytes were transfected with Lipofectamine LTX (Invitrogen, 15338030). Briefly, poly(I:C) was diluted in OPTI-MEM (1:100) and Lipofectamine LTX (1:2). The RNA-Lipofectamine solution incubated at room temperature for 25 minutes, then added dropwise to cells in OPTI-MEM and incubated at 37°C, 5% CO_2_.

### Viral infections

Vesicular Stomatitis Indiana Virus (VSV) and mutant for the M protein (VSV-M51) viruses were provided by David Olagnier (*45*). If not otherwise indicated, cells were infected with viruses at a Multiplicities Of Infection (MOI) of 10 and incubated for indicated time at 37°C, 5% CO_2_.

### Cell stimulation in 2D cultures

If not otherwise indicated, cells were plated at 3 × 10^4^, 5 × 10^4^, 7.5 × 10^4^, 1 × 10^5^ cells/well for 96-, 24-, 12-, and 6-well plates respectively, in complete culture medium. Next day, when indicated, cells were pre-incubated for 1 hour in OPTI-MEM (Gibco, 51985034) with inhibitors before the addition of toxins or control stimuli.

If not otherwise indicated, stimuli and inhibitors were used at the following concentrations: PLX-4720 (10 µM, MedChem Express, HY-51424); SB203580 (10 µM, MedChem Express, HY-10256); Doramapimod (10 µM, MedChem Express, HY-10320); BX795 (1 µM, Invivogen, Tlrl-bx7); Abemaciclib (25 nM, MedChem Express, HY-16297A); Ribociclib (40 nM, MedChem Express, HY-15777); Palbociclib (25 nM, MedChem Express, HY-50767); Fadraciclib (25 nM, MedChem Express, HY-101212); Roniciclib (25 nM, MedChem Express, HY-13914); SCH772984 (1 µM, MedChem Express, HY-50846); SP600125 (10 µM, MedChem Express, HY-12041); TAK-1 inhib HS-276 (10-20 µM, MedChem Express, HY-147141); TAO Kinase inhibitor 1 (5 µM, MedChem Express, HY-112136); Selonsertib (10 µM, MedChem Express, HY-18938); 5z-7-Oxozeaenol (5 µM, MedChem Express, HY-12686); GW806742X (1 µM, MedChem Express, HY-112292A); DN1289 (1 µM, MedChem Express, HY-152142); Gossypetin (40 µM, MedChem Express, HY-119917); BSJ-04-122 (10 µM, MedChem Express, HY-152185); GSK-872 (10 µM, MedChem Express, HY-101872); Bortezomib (1 µM, Selleck, SE-S1013-5MG); MLN9424 (1 µM, Tocris Bioscience, 6499); DT-061 (1-20 µM; MedChem Express, HY-112929); Dinophysistoxin (250 nM, Bertin Technologies); Okadaic acid (100-600 nM, Bertin Technologies, WP-10011490); Cantharidin (5-25 µM, Sigma-Aldrich, C7632-25MG); Calyculin-A (1 µM, Bertin Technologies, WP-19246); LB-100 (50 µM, Bertin Technologies, 29105); Raphin-1 (50 µM, Tocris Bioscience, 6760/10); Sephin-1 (100 µM, Tocris Bioscience, 5553/10); Anisomycin (1-5 µM, Selleck, SE-S7409-10MG); Valboro-Pro/ Talabostat (10 µM, Selleck, SE-S8455-5MG); Phosphatase Inhibitor Library (117 items) (10 µM, MedChem Express, HY-L081).

### ASC specks imaging and quantification

ASC specks formation in A549 and HEK293 cells was monitored using EVOS 7000 fluorescence microscope using x 10 or x 20 objectives. Specks were manually quantified as the proportion of ASC aggregates over total nuclei number (staining with Hoechst 33342, Invitrogen, H3570) using Fiji software (ImageJ). At least three wells and ≥500 cells per well were analysed in a blinded manner.

### Immunoblotting

Whole-cell lysates were prepared by lysing cells in RIP-A buffer (150 mM NaCl, 50 mM Tris-HCl, 1% Triton X-100, 0.5% Na-deoxycholate) supplemented with protease inhibitor cocktail (Roche, PHOSS-RO). For phosphorylated proteins samples, phosphatase inhibitor cocktail (Roche, 04693159001) was added to the lysis buffer. Before addition of 4X Laemmli buffer (Bio-Rad, 1610747) (with a 3: 1 ratio), protein concentration was measured with BCA protein assay (PIERCE, 23227). Lysates were then boiled at 95°C for 5 minutes.

Cell culture supernatants were precipitated by addition of TriChloroacetic Acid (TCA, 100% w/v, 4°C) to a final concentration of 10%, incubated on ice for 1 hour and centrifuged for 1 hour (15,000 rpm, 4°C). Pellets were washed with ice-cold acetone and centrifuged again (15,000 rpm, 4°C). The acetone was removed, proteins were resuspended in TE buffer and then boiled at 95°C for 5 minutes.

Samples were separated by home-made 12% denaturing SDS-PAGE, blotted onto 0.2-µm PVDF membranes (Bio-Rad, 12023927), and blocked in 5% milk diluted in TBS-T (Tris 10 mM, pH 8, NaCl 150 mM, Tween 20 0.05%). Membranes were incubated with indicated primary antibody overnight at 4°C and with corresponding secondary HRP-conjugated antibodies for 1 hour at room temperature. Chemiluminescent signals were detected with ECL revelation kit (Advansta, K-12043-D10, K-12042-D20) and recorded with C-DiGit Imaging System (Li-cor).

Antibodies used in this study were anti-Gasdermin D (E8G3F) Monoclonal antibody (1:1000, Cell signaling, 97558S); anti-Caspase 1 (p20) (human) mAb (Bally-1) antibody (1:500, Adipogen, AG-20B-0048); anti-DFNA5/Gasdermin E (EPR19859) Monoclonal antibody (1:1000,Abcam, ab215191); anti-Caspase 3 antibody (1:500, Cell signaling, 9662S); anti-Human IL-1β/IL-1F2 Polyclonal antibody (1:500, R&D systems, AF-201-NA); anti-Cleaved IL-1β (Asp116) Monoclonal antibody (1:750, Cell signaling, 83186S); anti-Tubulin-α antibody (1:1000, Abcam, ab4074); anti-PP1alpha Polyclonal antibody (1:1000, Invitrogen, PA5-119781); anti-PP1CB-Specific antibody (1:1000, ProteinTech, PR-55136-AP-150); anti-PP1CC antibody (1:1000, ProteinTech, PR-11082-1-AP-150); PP2A C Subunit antibody (1:500, Cell signaling, 2038S); anti-Purified NLRP1 (N-terminal) antibody (1:300, Biolegend, 679802); anti-NLRP1 (C-terminal) Polyclonal antibody (1:500,Abcam, ab36852); anti-GFP antibody (1:1000, Abcam, ab6673); anti-SNAP/CLIP-tag Monoclonal antibody (1:1000, ProteinTech, 6F9-100); anti-ZAKα Polyclonal antibody (1:1000, Bethyl Laboratories, A301-993A); anti-P38 MAPK antibody (1:1000, Cell signaling, 9212S); anti-Phospho-p38 (Thr180/Tyr182) (D3F9) Monoclonal antibody (1:1000, Cell signaling, 4511S); anti-Puromycine antibody Clone 12D10 (1:1000, Sigma-Aldrich, MABE343); anti-TAK1 antibody (1:1000, Cell signaling, 4505S); anti-Phospho-TAK1 (Ser412) antibody (1:1000, Cell signaling, 9339S); anti-Phospho-TAK1 (Thr184/187) 90C7 Monoclonal antibody (1:1000, Cell signaling, 4508S); anti-P38 alpha/MAPK14 antibody (E229) (1:1000, Abcam, ab170099); anti-P38 MAPK beta Monoclonal antibody (1:1000, Invitrogen, MA514-950); anti-P38 beta MAPK (C28C2) Monoclonal antibody (1:1000, Cell Signaling, 2339S); anti-P38 gamma/MAPK12 antibody (EPR6528N) (1:1000, Abcam, ab205926); anti-P38 gamma MAPK antibody (1:1000, Cell Signaling, 2307S); anti-Human P38 delta antibody (1:1000, R&D system, AF1519); anti-P38δ MAPK (10A8) mAb (1:1000, Cell Signaling, 2308S); anti-GAPDH antibody (1:1000, GeneTex, GTX100118); Goat anti-rabbit HRP secondary antibody (1:5000, Advansta R-05072-500); Goat anti-mouse HRP secondary antibody (1:5000, Advansta, R-05071-500); Goat anti-rat IgG H&L (HRP) (1:5000, Abcam, ab97057); Goat IgG HRP-conjugated Antibody (1:5000, Biotechne, HAF109).

### PhosTag SDS-PAGE

Cells were lysed in RIP-A buffer supplemented with protease inhibitor and phosphatase inhibitor cocktails, and processed as described above. Phosphorylated proteins were separated by home-made 10% SDS-PAGE gel, with addition of Phos-tag Acrylamide (Wako Chemicals, AAL-107) to a final concentration of 30 µM and Manganese Chloride (II) (Sigma-Aldrich, 63535) to 60 µM. Gel were washed in Transfer buffer containing EDTA (10 mM), blotted onto 0.45-µm PVDF membranes (Invitrogen, LV2005), and blocked in 5%. Membranes were incubated with indicated primary antibody and corresponding secondary HRP-conjugated antibodies.

### Plasma membrane permeabilization monitoring

N/TERT cells were seeded at 2 × 10^4^ cells/well in Black/Clear 96-well plates (Greiner, 655090) in complete culture medium. Twenty-four hours later, culture medium was changed for OPTI-MEM supplemented with SYTOX-Green Nucleic Acid Stain (500 nM, Invitrogen, S7020) and treated with the indicated stimulations. Real-time fluorescence was measured using Clariostar (BMG Labtech) plate reader equipped with a 37°C incubator. Maximal permeabilization was defined using 1% Triton X-100.

### Cell lysis assay

Cell lysis was evaluated by the quantification of LDH release into the cell supernatant, employing the LDH-Blue kit (rep-ldh-1) from Invivogen. Keratinocytes were seeded in 96-well plates at 2 × 10^4^ cells per well in Keratinocyte Growth Medium 2 (PromoCell Inc). The following day, cells were stimulated and 50 µL of cell supernatant was mixed with an equal volume of LDH substrate and left to incubate for 30 min at room temperature, protected from light. The enzymatic reaction was stopped by adding 50 µL of stop solution. Maximal cell death was determined with whole cell lysates from unstimulated cells incubated with 1% Triton X-100.

### Kinase activity assay

Kinase activity profiling was performed using PamChip 96-microarrays (PamGene, 86311). N/TERT keratinocytes were stimulated with dinophysistoxin or not for 3 hours, washed in ice-cold PBS, and lysed in manufacturer-recommended lysis buffer. Protein concentrations were normalised using BCA assay and equal amounts of lysates were loaded per array, according to the supplier’s protocol. Fluorescent signals were acquired and quantified in BioNavigator software. Technical replicates passing automated quality control were averaged to generate one kinase activity profile per biological replicate (n = 3 per condition). Differential kinase activity was determined by comparing dinophysistoxin-treated samples to matched controls.

### Cytokine analysis / quantification / Soluble mediator release analysis

To measure hIL-1β levels, cell supernatants were recovered 24 hours after stimulation and human IL-1β Enzyme Linked Immunosorbent Assay (ELISA) kit (Invitrogen, 88-7261-77) was used according to the supplier’s protocol.

For the Human IL-1 Family Cytokine Array (Tebu-bio, AAH-IL1F-1-8), N/TERT were seeded in T-75 flask at 5 × 10^5^ cells per wells in complete culture medium. Supernatants were collected 24 hours after stimulation, cleared by centrifugation (15 000 rpm, 10 minutes, 4°C), concentrated with Amicon Ultra 4 – 10K (4,000 g, 15 minutes, 4°C, Sigma-Aldrich, UFC801024), and cytokine array was performed according to the supplier’s protocol.

To measure secreted cytokine and soluble mediators’ levels for human skin models, culture mediums were collected, cleared by centrifugation (15 000 rpm, 10 minutes, 4°C), and LEGENDPlex^TM^ Human cytokine panel 2 (Biolegend, 741378) was used according to the supplier’s protocol. Samples were processed with BD Fortessa x20 Nothern Lights C6 L2 (Genotoul Tri, IPBS, toulouse) and analysed using the LEGENDplex Data Analysis Software Suite (Biolegend). The following targets were analysed: TSLP, IL-1α, IL-1β, GM-CSF, IFN-α2, IL-23, IL-12p40, IL-12p70, IL-15, IL-18, IL-11, IL-27, and IL-33

### Histology

Human skin tissue samples were fixed in a 10% formalin bath for 48 hours at room temperature, dehydrated automatically and embedded in paraffin (Epredia HistoStar, Epredia). Paraffin blocks were cut into 5 µm-thick sections and stained with Hematoxylin and Eosin (H&E) according to standard protocols. Images were acquired using a Zeiss Axio Imager M2 (Zeiss) and 20x/0.8 EC Plan-neofluar Zeiss objective (Zeiss). Images were acquired and processed using Zeiss Axiocam 503 color camera and the Zeiss Zen software (Zeiss).

### Immunohistology

Human skin tissue samples were snap-frozen in liquid nitrogen and embedded in Tissue Freezing Medium (TFM) (Microm Microtech France, TFM-5). Frozen blocks were cut into 7 µm-thick sections (Phi plateform, I2MC, Toulouse), fixed in acetone 100% for 20 minutes at room temperature and rehydrated in PBS for 15 minutes. Sections were then blocked for 1 hour at room temperature with 3% Bovine Serum Albumin in PBS-T (PBS 0.1% Tween 20) and incubated overnight at 4°C in a humidity chamber with primary antibody (Cytokeratin K-14,1:400, Abcam, Ab7800; ASC,1:250, AdipoGen, AG-25B-0006) diluted in staining buffer (1% BSA, PBS 0.05% Tween). Slides were then incubated with Dylight 550 (1:500, ImmunoReagents Inc., GtxMu-003-D550NHSX), Dylight 488 (1:500, ImmunoReagents Inc., GtxRb-003-0488NHSX) -Conjugated secondary antibodies in staining buffer for 1 hour at room temperature. Hoechst (1:1000) was added to the last wash for 10 minutes at room temperature. Wild-field images were acquired using EVOS 7000 fluorescence microscope with x 20 objectives and confocal imaging was performed on a Spinning disk Andor (Olympus) with 60XO/1.35 ULSAPO (WD 0.15mm) objective at the Genotoul Imagerie plateform (IPBS, Toulouse). Images were processed using IQ3 software (Olympus).

### Proximity Ligation assay

ASC-NLRP1 interactions were visualised on frozen sections (as described above) using the NaveniFlex Tissue MR (Navinci, 60027) kit, according to manufacturer’s protocol. Briefly, sections were blocked for 1 hour with Blocking buffer, incubated overnight at 4°C with primary antibody NLRP1 (1:200, Biolegend, 679802), ASC (1:200, AdipoGen, AG-25B-0006) and incubated with Navenibody M1 and Navenibody R2 diluted in Diluent buffer (1:40) for 1 hour. Slides were incubated with Enzyme 1 diluted in 1X Buffer 1 (1:40) for 30 minutes and with Enzyme 2 (1:40) for 90 minutes. For visualisation, slides were incubated in Detection buffer for 30 minutes, incubated in PBS supplemented with Hoechst (1:1000) for 5 minutes and mounted in mounting medium. Otherwise specified, every incubation step was made at 37°C in a humidity chamber. Images were acquired using Confocal and Multiphoton Zeiss 710 NLO Spectral (Zeiss) and 20X/0.8 PLAN APO AIR objective. Images were processed using Zeiss Zen software (Zeiss).

### Quantification of epithelial damage and ASC specks in 3D skin models

Intradermal disruptions were quantified as the ratio of disruption area to total epidermis area. Regions of interest were manually delineated in Fiji using H&E sections from three technical replicates per treatment within the same experiment.

ASC specks and Proximity Ligation Assay (PLA)-positive puncta in tissue sections were quantified as the proportion of ASC aggregates or PLA signals per nucleus using Fiji software. Quantifications were performed on images from three technical replicates per treatment within the same experiment.

### Proteins expression and purification

HEK 293T cells were seeded at 3 × 10^5^ cells per well on a 6-well plate and transfected with a plasmid coding for NLRP1 Disordered Region (DR) construct (aa 86-254) expressing a GFP tag (*22*). Transfection was made using Lipofectamine 2000 (Invitrogen, 11668027) and 2 µg of DNA per well, according to manufacturer’s protocol. Cells were lysed in RIP-A buffer and cleared by centrifugation (17 000 g, 10 minutes, 4°C). Immunoprecipitation (IP) was conducted using GFP-Trap Agarose (Chromotek, gta) kit, according to supplier’s protocol. Briefly, diluted lysates were added to equilibrated beads (in dilution buffer) and incubated for 1 hour at 4°C on a rotating wheel. Subsequently, beads were washed four times with Wash buffer before elution in homemade acidic elution buffer (0.1 M Glycine pH 2.7) and neutralisation with 1 M Tris pH 8.5. Purified NLRP1-DR-GFP was quantified using Bradford/ BCA assay and used for lambda phosphatase dephosphorylation assay.

### Lambda Phosphatase dephosphorylation assay

Purified hNLRP-GFP-DR was adjusted to a final volume of 40 µL with distilled water, yielding at 40 µg protein per reaction. Samples were supplemented with 10X NEBuffer for Protein MetalloPhosphatases (PMP, 5 µL), MnCl_2_ (10 mM, 5 µL), and Lambda Protein Phosphatases (1 µL, New England Biolab, P0753S). Reactions were incubated at 30°C for 30 minutes. Following dephosphorylation, samples were processed for SDS-PAGE phostag immunoblotting as described previously in the Phostag SDS-PAGE section.

### *In vitro* kinase assay

Kinase reactions were carried out by combining 500 ng of recombinant NLRP1 (Clinisciences, P316481) with 100 ng of either recombinant p38α (Abcam, ab271606) or TAK1 (Abcam, ab89692) in homemade reaction buffer (HEPES-KOH pH 7.0, 5 mM MgCl_2_, 2 mM ATP, 0.3 M DTT) for 1 hour at 30°C. When indicated, kinase products were subsequently exposed to Lambda Protein Phosphatase in 1X NEBuffer for PMP and MnCl_2_ (1 mM). Mixtures were incubated for 30 minutes at 30°C. Final reaction products were processed for SDS-PAGE Phostag immunoblotting as described previously in the Phostag SDS-PAGE section.

### Statistical analysis

Statistical analyses were performed using GraphPad Prism (version 10, GraphPad Software, Inc.). The exact test, sample size (n), and correction methods are provided in the corresponding figure legend. Unless otherwise specified, data are presented as mean ± SEM. Comparisons between two or more groups were analysed using one-way ANOVA or two-way ANOVA (respectively) with the appropriate multiple-comparison correction. P values are reported in the figures, with the following significance thresholds: *P ≤ 0.05, **P ≤ 0.01, ***P ≤ 0.001; ns indicates no significant difference.

## Acknowledgments

The authors acknowledge all lab members as well as members from FL Zhong’s and P. Mitchell’s labs from fruitful discussions. for their critical advices and support on this project. The authors also acknowledge the IPBS microscopy, cytometry and histology platforms from IPBS and I2MC institutes.

## Author contribution

Conceptualization: MP, EM

Methodology: MP, LG, NG, FLZ, DO, EM

Investigation: MP, LG, LSDC, RC, GA, AM, LB, TB, AG, CB, LRK, BS, LF, RSO, DP, EM

Visualization: MP, LG, DO, EM, FLZ, NG, VS, LB, CC

Funding acquisition: EM, CC, DO

Project administration: EM Supervision: EM, RM, CC, DO

Writing – original draft: MP, LG, EM

Writing – review & editing: MP, LG, EM, DO, RM

## Competing interests

Authors declare that they have no competing interests.

All data are available in the main text or the supplementary materials.

All tools generated in this study are available upon request to

## Supplemental information

### Supplemental Figures

**Supplemental Figure 1 (Refers to Figure 1).**
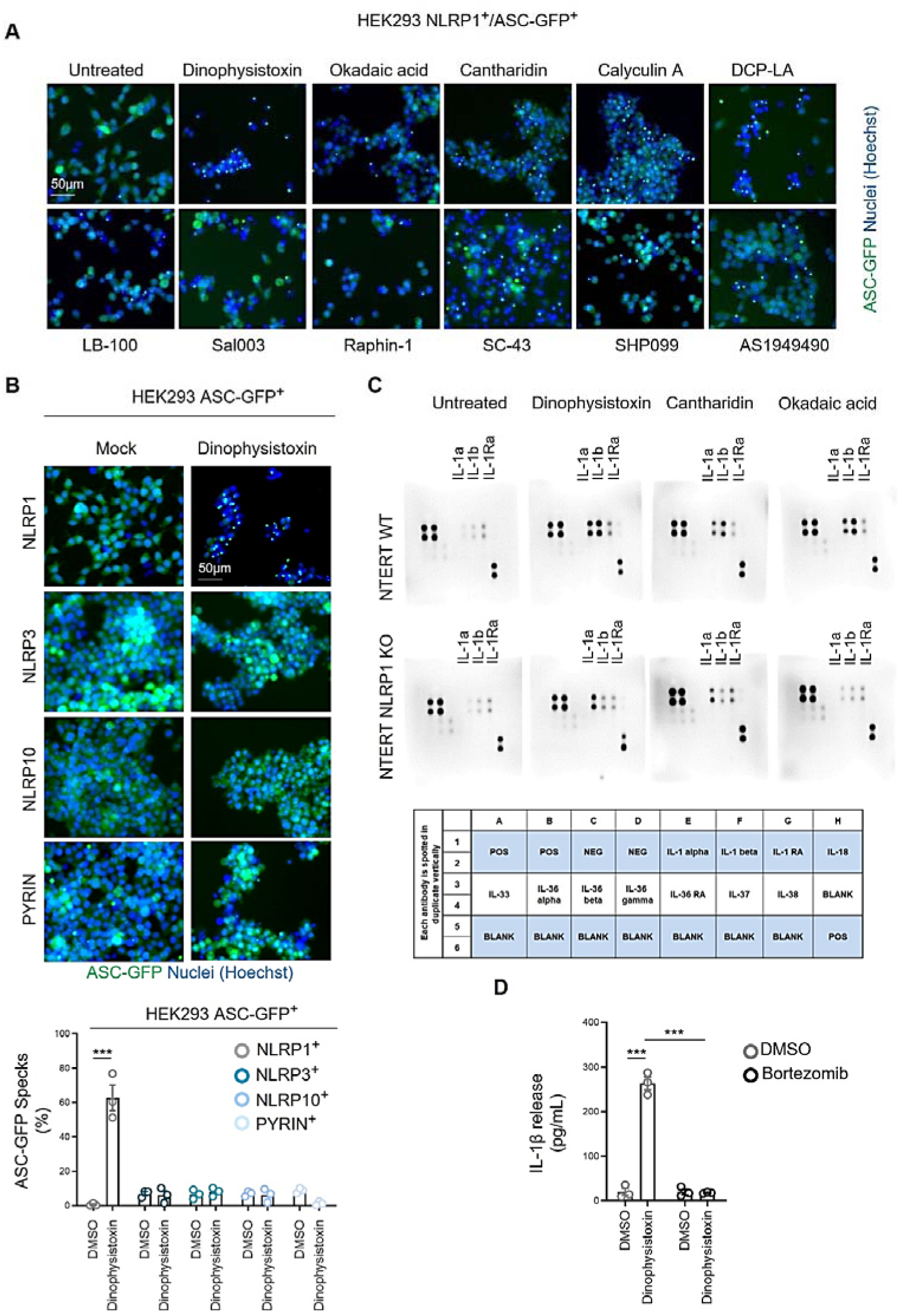
Multiple PP1/PP2A inhibitors trigger the human NLRP1 inflammasome activation. **A.** Fluorescence microscopy of ASC-GFP specks in WT HEK293T^ASC-GFP/NLRP1^ reporter cells exposed to selected phosphatase inhibitory compounds for 8 hours. ASC-GFP (green) pictures were directly taken in dish after adding Hoechst (nuclei staining). Images shown are from one experiment and are representative of three independent experiments; scale bars, 50 µm. ASC complex percentage was performed by determining the ratios of cells positive for ASC speckles on the total nuclei (Hoechst). **B.** Fluorescence microscopy of ASC-GFP specks in multiple HEK293T^ASC-GFP^ reporter cells expressing NLRP1, NLRP10, PYRIN or NLRP3 inflammasome-forming sensors and exposed to selected Dinophysistoxin (100nM) compounds for 8 hours. ASC-GFP (green) pictures were directly taken in dish after adding Hoechst (nuclei staining). Images shown are from one experiment and are representative of three independent experiments; scale bars, 50 µm. ASC complex percentage was performed by determining the ratios of cells positive for ASC speckles on the total nuclei (Hoechst). **C.** Cytokine analysis 8 h after exposure of WT and NLRP1KO TERT keratinocytes to Dinophysistoxin (100nM), Okadaic acid (250nM) and Cantharidin (5µM). Representative experiment of three independent replicates. **D.** Determination of IL-1β release in WT NTERT-keratinocytes after 8 h exposure to Dinophysis toxin (Dino. toxin, 100nM) in presence/absence of bortezomib (1µM). ***P ≤ 0.0001, one-way ANOVA. Values are expressed as mean ± SEM. Graphs show one experiment performed in triplicates at least three times.

**Supplemental Figure 2 (Refers to Figure 4).**
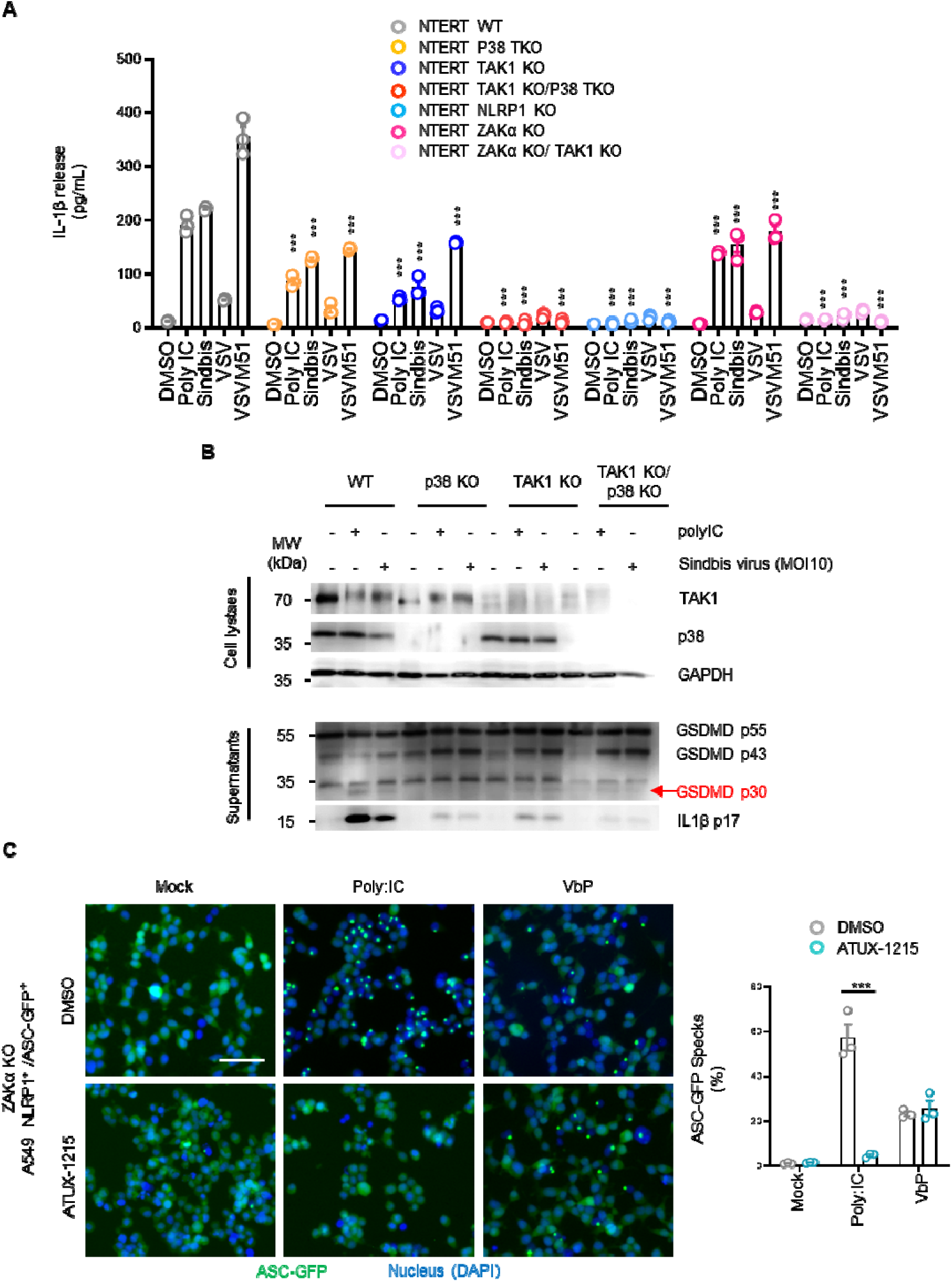
TAK1-driven NLRP1 inflammasome activation expands to dsRNA and viral infections. **A.** Determination of IL-1β release in WT, P38 TKO, TAK1 KO, NRLP1 KO, ZAKα KO, TAK1 KO/ P38 TKO and ZAKα KO/ TAK1 KO NTERT-keratinocytes after 24 h infection Sindbis, VSV and VSVM51 viruses (MOI 10) or after Poly:IC (5µg/mL) transfection. ***P ≤ 0.0001, one-way ANOVA. Comparisons of each treatment in various genotype to it respective WT condition. Values are expressed as mean ± SEM. Graphs show one experiment performed in triplicates at least three times. **B.** Immunoblotting of P38, TAK1, cleaved GSDMD and IL-1β in WT, P38α/β/δ TKO, TAK1 KO, P38α/β/δ TKO/TAK1 KO in NTERT keratinocytes infected or not with Sindbis virus (MOI 10) for 30 hours or transfected with poly:IC (5µg/mL). Pictures show one experiment performed at least two times. **C.** Fluorescence microscopy and associated quantifications of ASC-GFP specks in ZAKα KO A549^ASC-GFP/NLRP1^ reporter cells exposed to VbP (5µM) or transfected with poly:IC (5µg/mL) for 6 hours in presence/absence of the PP2A activator ATUX-1215 (15µM). ASC-GFP (green) pictures were directly taken in dish after adding Hoechst (nuclei staining). Images shown are from one experiment and are representative of three independent experiments; scale bars, 50 µm. ASC complex percentage was performed by determining the ratios of cells positive for ASC speckles on the total nuclei (Hoechst).

**Supplemental Figure 3 (Refers to Figure 5).**
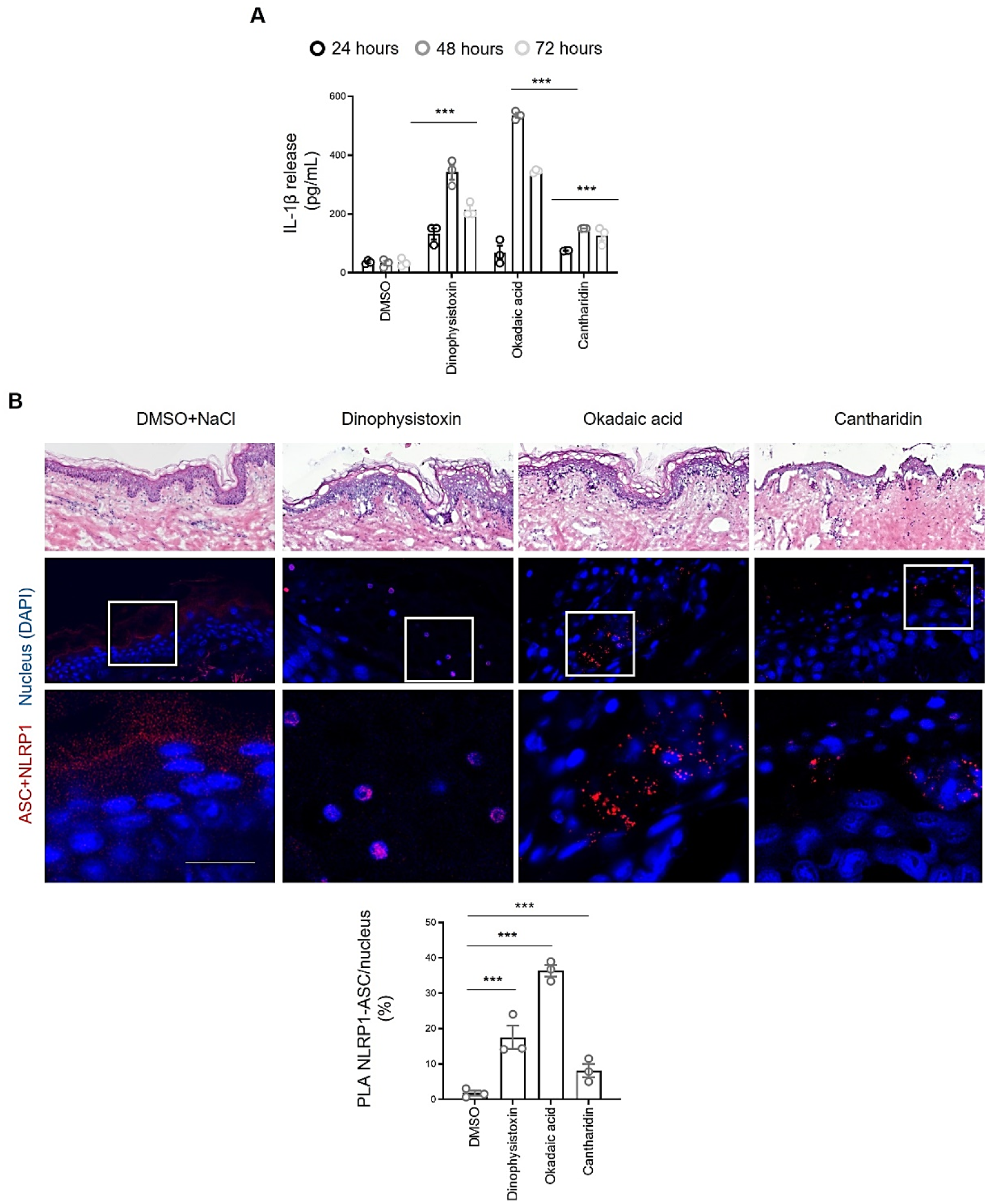
Extended results regarding Human native skins. **A.** Evaluation of IL-1β release upon exposure of Human skin explants to Dinophysistoxin (250 nM), Okadaic acid (600 nM) and Cantharidin (10µM) for various time. ***P ≤ 0.0001, one-way ANOVA compared to their respective PBS controls. Values are expressed as mean ± SEM. Graphs show one experiment performed in duplicate at least two times. **B.** Hematoxylin (H) & Eosin (E), ASC-NLRP1 Proximity Ligation Assay (PLA) staining and associated quantifications showing ASC and NLRP1 in close proximity after exposure of Human native skins to Dinophysis toxin (250 nM), Okadaic acid (600 nM) and Cantharidin (10µM) for 24 hours. P values indicated in figure, one-way ANOVA. Images are representative of three biological replicates. Scale bar = 50 µm.

## Supplemental material

**Table S1.**
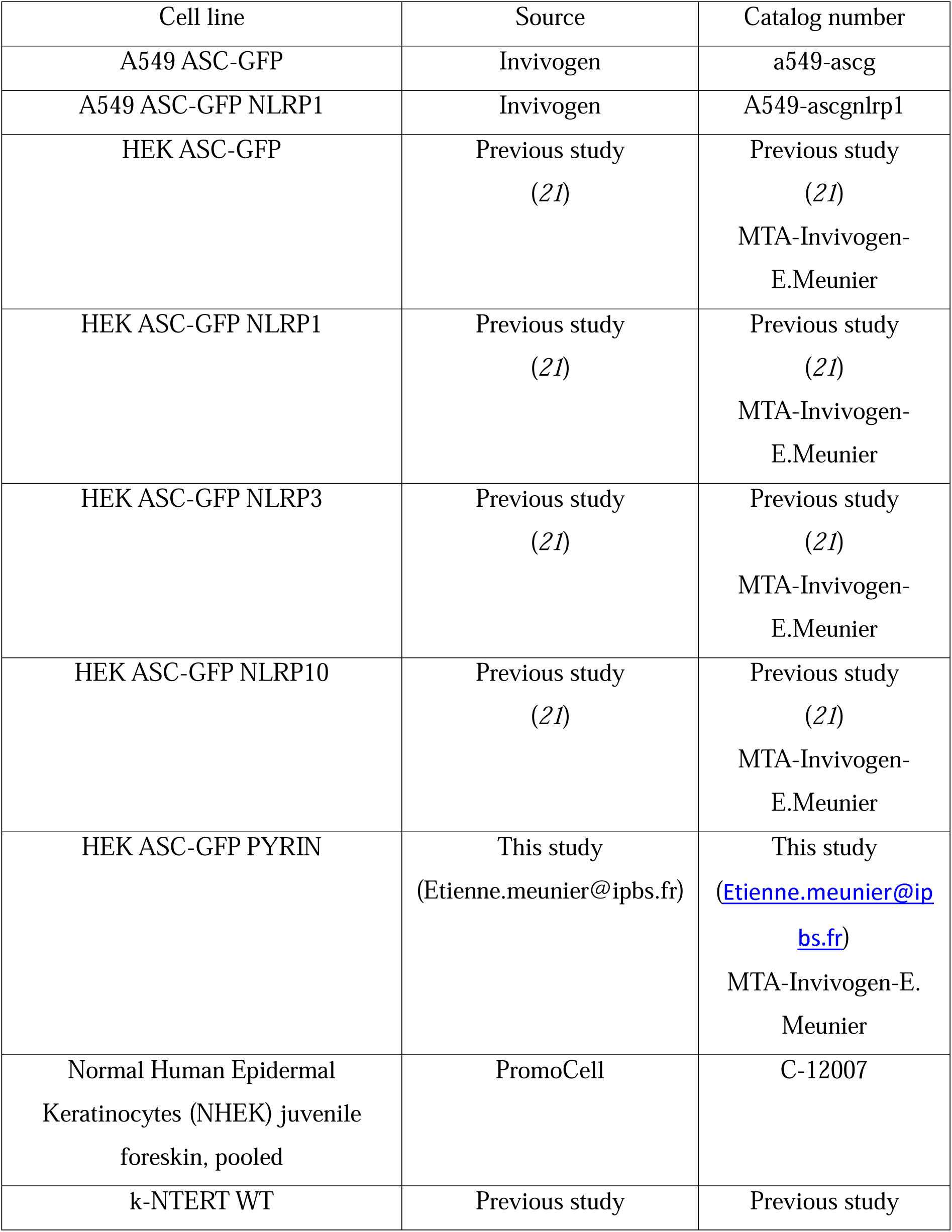
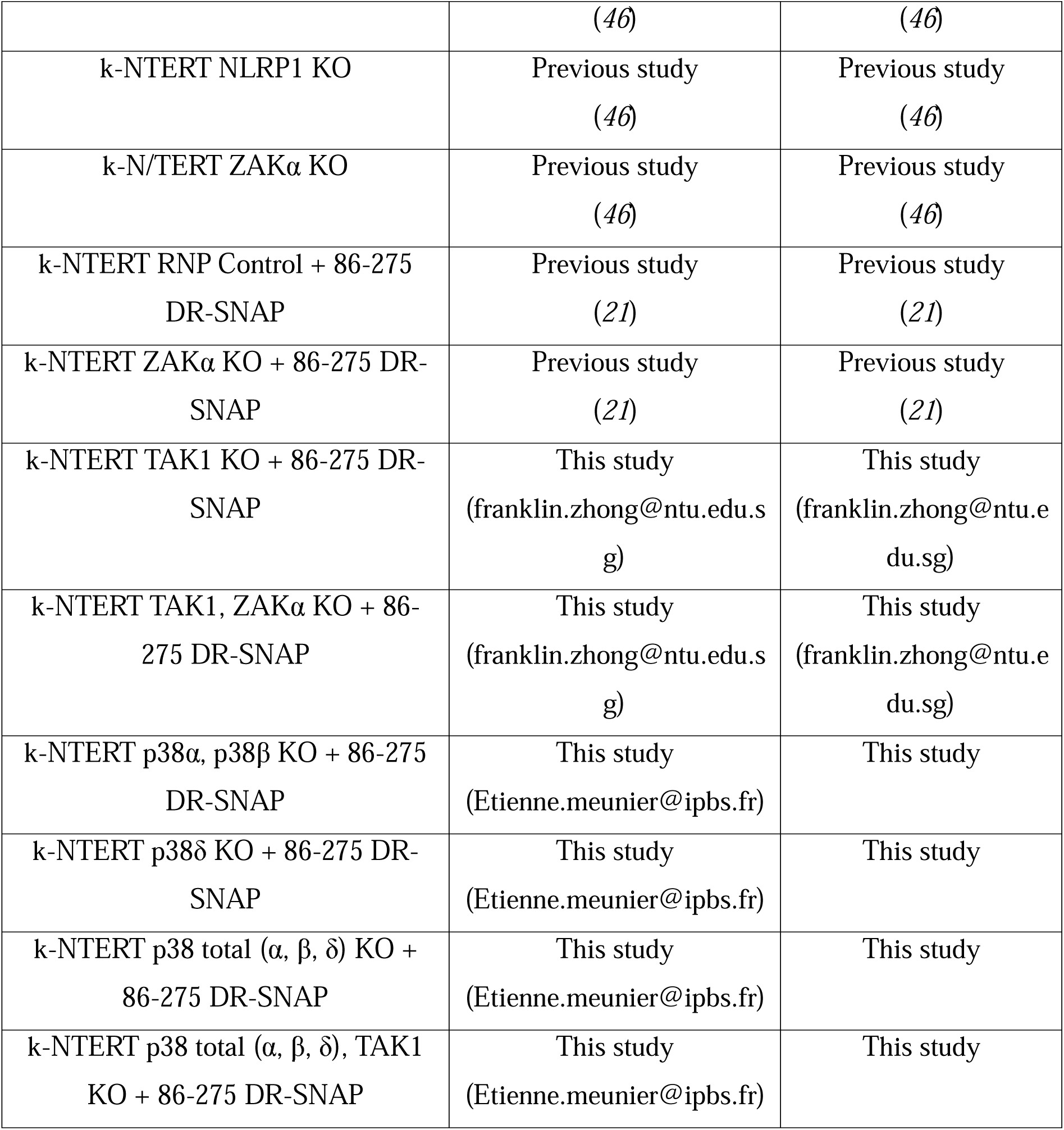
List of all cell lines used in this study is provided in Table S1.

