## Supplemental Material for "A TAK1-Driven NLRP1 Inflammasome Pathway Revealed by Phosphatase-Targeting Environmental Toxins"

**Abstract**

Human NLRP1 is highly sensitive to phosphorylation-dependent activation, indicating a need for tight phosphatase control to prevent excessive inflammasome activity. In this study, we identified the phosphatases PP1 and PP2A as negative regulators of the NLRP1 inflammasome in human keratinocytes. Accordingly, exposure to the environmental toxins Dinophysistoxin, Okadaic acid, and Cantharidin, which inhibit PP1 and PP2A, triggered NLRP1 inflammasome activation. Notably, this toxin-induced activation process relied on hyperactivation of the MAP3 kinase TAK1. Mechanistically, both TAK1 and its downstream effectors p38 kinases phosphorylated and activated the NLRP1 inflammasome. Further studies underscored that TAK1 also contributed to the NLRP1 inflammasome activation during double-stranded RNA stimulation and viral infection. Finally, human native skins exposed to environmental toxins underlined the role of the PP1/PP2A-regulated TAK1/p38 axis in the development of skin dermatitis. Thus, these findings reveal a novel pathway of phosphorylation-driven NLRP1 activation and expand our understanding of its regulation in epithelial immunity.

**One Sentence Summary:** PP1/PP2A phosphatases restrict TAK1-driven NLRP1 inflammasome response.

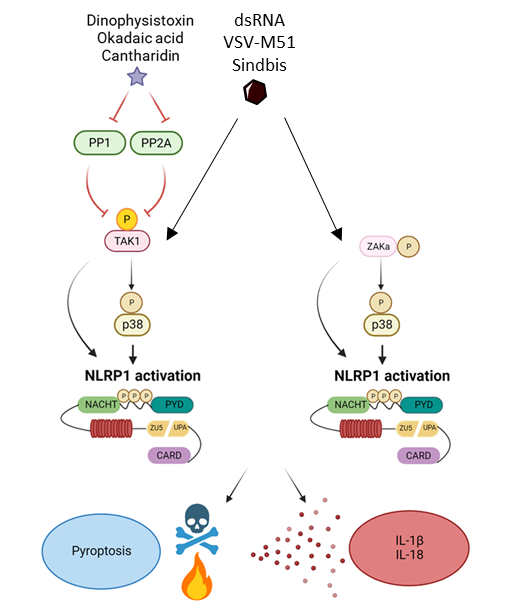

**Graphical abstract.** Schematic representation of the phosphorylation-driven hNLRP1 inflammasome response upon exposure to environmental toxins and to dsRNA/viral infection.

**Supplemental information**

**Supplemental Figures**

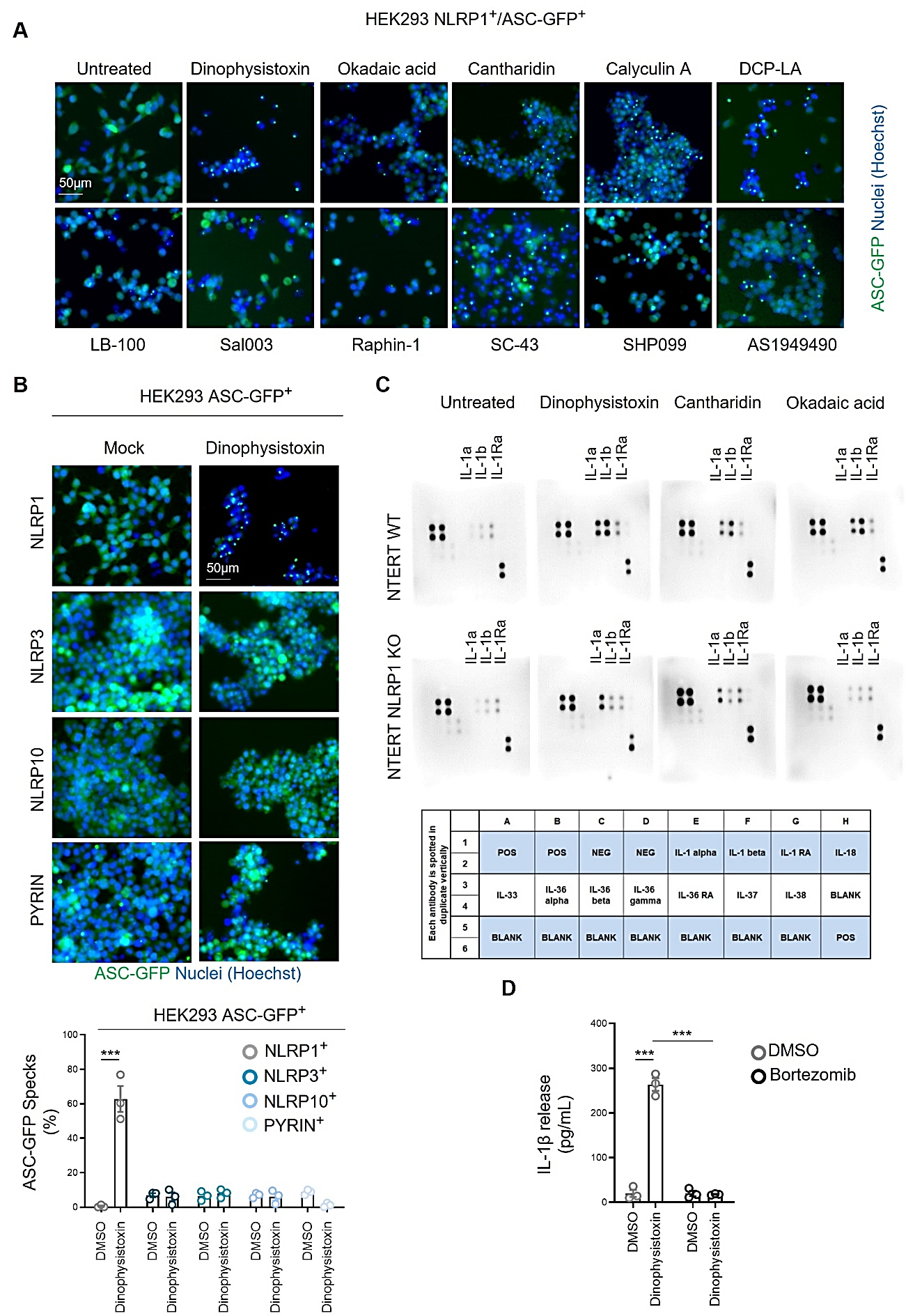

**Supplemental Figure 1 (Refers to Figure 1). Multiple PP1/PP2A inhibitors trigger the human NLRP1 inflammasome activation**

**A.** Fluorescence microscopy of ASC-GFP specks in WT HEK293T^ASC-GFP/NLRP1^ reporter cells exposed to selected phosphatase inhibitory compounds for 8 hours. ASC-GFP (green) pictures were directly taken in dish after adding Hoechst (nuclei staining). Images shown are from one experiment and are representative of three independent experiments; scale bars, 50 µm. ASC complex percentage was performed by determining the ratios of cells positive for ASC speckles on the total nuclei (Hoechst).

**B.** Fluorescence microscopy of ASC-GFP specks in multiple HEK293T^ASC-GFP^ reporter cells expressing NLRP1, NLRP10, PYRIN or NLRP3 inflammasome-forming sensors and exposed to selected Dinophysistoxin (100nM) compounds for 8 hours. ASC-GFP (green) pictures were directly taken in dish after adding Hoechst (nuclei staining). Images shown are from one experiment and are representative of three independent experiments; scale bars, 50 µm. ASC complex percentage was performed by determining the ratios of cells positive for ASC speckles on the total nuclei (Hoechst).

**C.** Cytokine analysis 8 h after exposure of WT and NLRP1KO TERT keratinocytes to Dinophysistoxin (100nM), Okadaic acid (250nM) and Cantharidin (5µM). Representative experiment of three independent replicates.

**D.** Determination of IL-1β release in WT NTERT-keratinocytes after 8 h exposure to Dinophysis toxin (Dino. toxin, 100nM) in presence/absence of bortezomib (1µM). ***P ≤ 0.0001, one-way ANOVA. Values are expressed as mean ± SEM. Graphs show one experiment performed in triplicates at least three times.

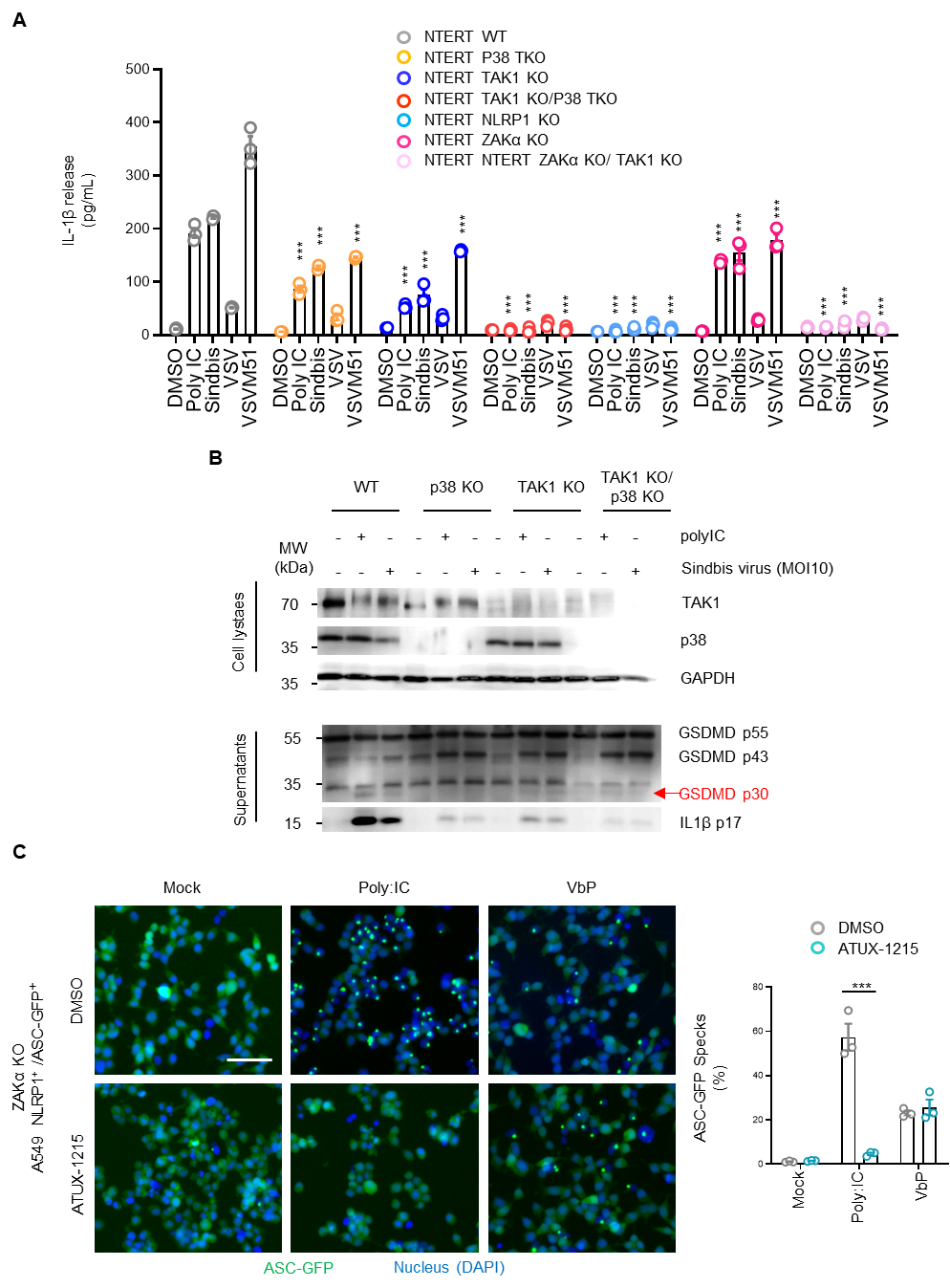

**Supplemental Figure 2 (Refers to Figure 4). TAK1-driven NLRP1 inflammasome activation expands to dsRNA and viral infections**

**A.** Determination of IL-1β release in WT, P38 TKO, TAK1 KO, NRLP1 KO, ZAKα KO, TAK1 KO/ P38 TKO and ZAKα KO/ TAK1 KO NTERT-keratinocytes after 24 h infection Sindbis, VSV and VSVM51 viruses (MOI 10) or after Poly:IC (5µg/mL) transfection. ***P ≤ 0.0001, one-way ANOVA. Comparisons of each treatment in various genotype to it respective WT condition. Values are expressed as mean ± SEM. Graphs show one experiment performed in triplicates at least three times.

**B.** Immunoblotting of P38, TAK1, cleaved GSDMD and IL-1β in WT, P38α/β/δ TKO, TAK1 KO, P38α/β/δ TKO/TAK1 KO in NTERT keratinocytes infected or not with Sindbis virus (MOI 10) for 30 hours or transfected with poly:IC (5µg/mL). Pictures show one experiment performed at least two times.

**C.** Fluorescence microscopy and associated quantifications of ASC-GFP specks in ZAKα KO A549^ASC-GFP/NLRP1^ reporter cells exposed to VbP (5µM) or transfected with poly:IC (5µg/mL) for 6 hours in presence/absence of the PP2A activator ATUX-1215 (15µM). ASC-GFP (green) pictures were directly taken in dish after adding Hoechst (nuclei staining). Images shown are from one experiment and are representative of three independent experiments; scale bars, 50 µm. ASC complex percentage was performed by determining the ratios of cells positive for ASC speckles on the total nuclei (Hoechst).

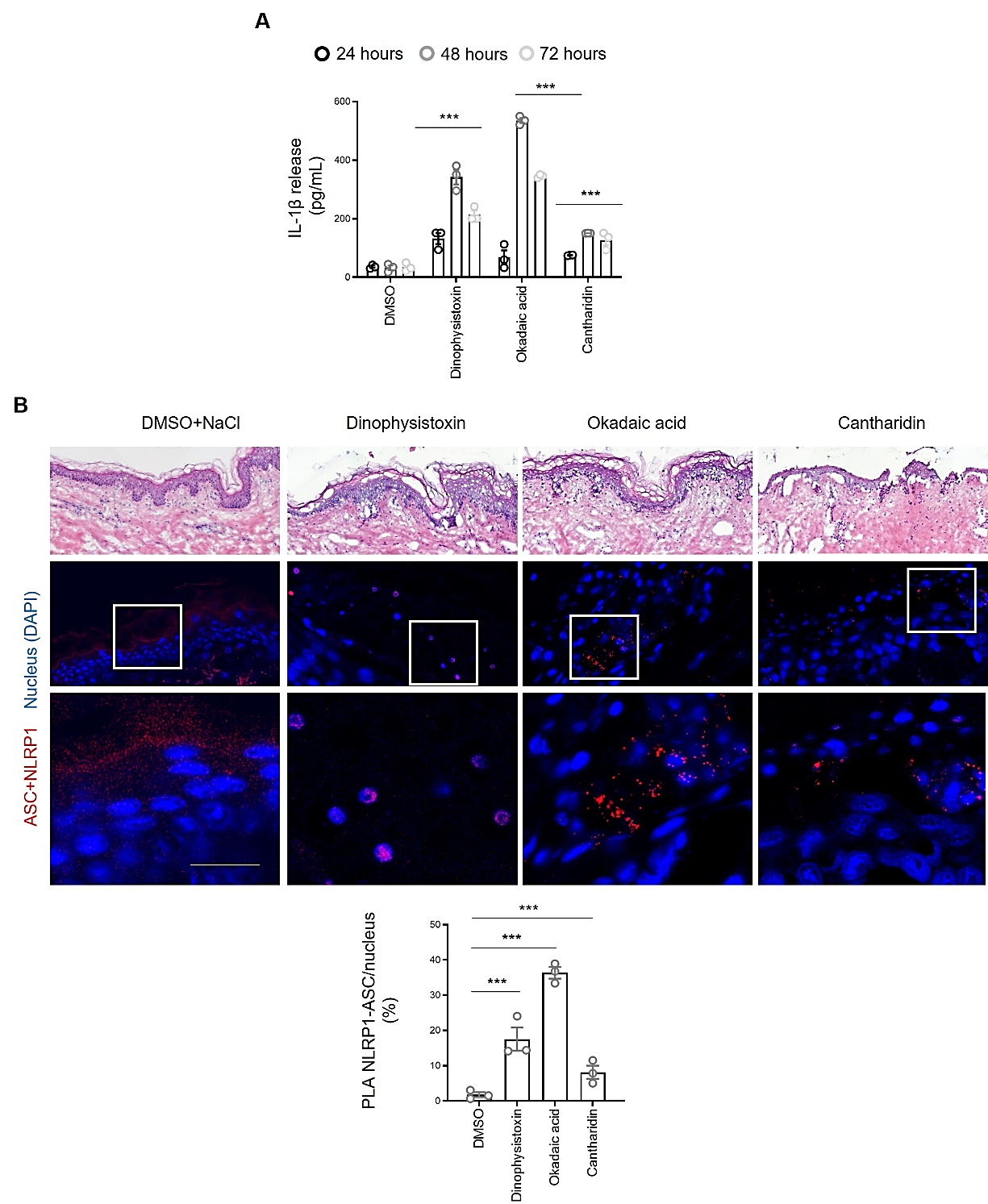

**Supplemental Figure 3 (Refers to Figure 5). Extended results regarding Human native skins**

**A.** Evaluation of IL-1β release upon exposure of Human skin explants to Dinophysistoxin (250 nM), Okadaic acid (600 nM) and Cantharidin (10µM) for various time. ***P ≤ 0.0001, one-way ANOVA compared to their respective PBS controls. Values are expressed as mean ± SEM. Graphs show one experiment performed in duplicate at least two times.

**B.** Hematoxylin (H) & Eosin (E), ASC-NLRP1 Proximity Ligation Assay (PLA) staining and associated quantifications showing ASC and NLRP1 in close proximity after exposure of Human native skins to Dinophysis toxin (250 nM), Okadaic acid (600 nM) and Cantharidin (10µM) for 24 hours. P values indicated in figure, one-way ANOVA. Images are representative of three biological replicates. Scale bar = 50 µm.

**Supplemental material**

**Table S1**. List of all cell lines used in this study is provided in **Table S1**.

| Cell line | Source | Catalog number |
| --- | --- | --- |
| A549 ASC-GFP | Invivogen | a549-ascg |
| A549 ASC-GFP NLRP1 | Invivogen | A549-ascgnlrp1 |
| HEK ASC-GFP | Previous study  (*21*) | Previous study  (*21*)  MTA-Invivogen-E.Meunier |
| HEK ASC-GFP NLRP1 | Previous study  (*21*) | Previous study  (*21*)  MTA-Invivogen-E.Meunier |
| HEK ASC-GFP NLRP3 | Previous study  (*21*) | Previous study  (*21*)  MTA-Invivogen-E.Meunier |
| HEK ASC-GFP NLRP10 | Previous study  (*21*) | Previous study  (*21*)  MTA-Invivogen-E.Meunier |
| HEK ASC-GFP PYRIN | This study  | This study   MTA-Invivogen-E. Meunier |
| Normal Human Epidermal Keratinocytes (NHEK) juvenile foreskin, pooled | PromoCell | C-12007 |
| k-NTERT WT | Previous study  (*45*) | Previous study  (*45*) |
| k-NTERT NLRP1 KO | Previous study  (*45*) | Previous study  (*45*) |
| k-N/TERT ZAKα KO | Previous study  (*45*) | Previous study  (*45*) |
| k-NTERT RNP Control + 86-275 DR-SNAP | Previous study  (*21*) | Previous study  (*21*) |
| k-NTERT ZAKα KO + 86-275 DR-SNAP | Previous study  (*21*) | Previous study  (*21*) |
| k-NTERT TAK1 KO + 86-275 DR-SNAP | This study  | This study  |
| k-NTERT TAK1, ZAKα KO + 86-275 DR-SNAP | This study  | This study  |
| k-NTERT p38α, p38β KO + 86-275 DR-SNAP | This study  | This study |
| k-NTERT p38δ KO + 86-275 DR-SNAP | This study  | This study |
| k-NTERT p38 total (α, β, δ) KO + 86-275 DR-SNAP | This study  | This study |
| k-NTERT p38 total (α, β, δ), TAK1 KO + 86-275 DR-SNAP | This study  | This study |
